# CryoSIM: super resolution 3D structured illumination cryogenic fluorescence microscopy for correlated ultra-structural imaging

**DOI:** 10.1101/2020.03.09.980334

**Authors:** Michael A. Phillips, Maria Harkiolaki, David Miguel Susano Pinto, Richard M. Parton, Ana Palanca, Manuel Garcia-Moreno, Ilias Kounatidis, John W. Sedat, David I. Stuart, Alfredo Castello, Martin J. Booth, Ilan Davis, Ian M. Dobbie

**Affiliations:** Micron Advanced Bio-imaging Unit, Department of Biochemistry, University of Oxford, South Parks rd, Oxford OX1 3QU, UK; STRUBI, Division of Structural Biology, Wellcome Centre for Human Genetics, Old Road Campus, Roosevelt Drive, Oxford, OX3 7BN, UK; Diamond Light Source, Harwell Science & Innovation Campus, Didcot, Oxfordshire, OX11 0DE, UK; Department of Anatomy and Cell Biology, Faculty of Medicine, Universidad de Cantabria, CP39011 Santander, Spain; Department of Biochemistry, University of Oxford, South Parks rd, Oxford OX1 3QU, UK; Department of Biochemistry and Biophysics, University of California, San Francisco, USA; Department of Engineering Science, University of Oxford, Parks Road, Oxford,OX1 3PJ, UK

## Abstract

Rapid cryo-preservation of biological specimens is the gold standard for visualising cellular structures in their true structural context. However, current commercial cryo-fluorescence microscopes are limited to low resolutions. To fill this gap, we have developed cryoSIM, a microscope for 3D super-resolution fluorescence cryo-imaging for correlation with cryo electron microscopy or cryo soft X-ray tomography. We provide the full instructions for replicating the instrument mostly from off-the-shelf components and accessible, user-friendly open source Python control software. Therefore, cryoSIM democratises the ability to detect molecules using super-resolution fluorescence imaging of cryo-preserved specimens for correlation with their cellular ultrastructure.

## 1. Introduction

Imaging methods for cells and tissues have progressed rapidly in the past decade, providing unrivalled opportunities for new insights into biological molecular mechanisms [1–3]. While many of the highest profile developments have been in super-resolution fluorescence and single molecule methods, other innovations in electron or X-ray microscopy have facilitated visualisation of ultrastructural morphology [4, 5]. The ultrastructural information becomes particularly informative when the distributions of specific molecules are added. This can be done by imaging the same sample in fluorescence and precisely painting the fluorescence signal onto the EM or X-ray tomography deduced morphology in so called correlative methods, producing correlative light and electron microscopy (CLEM) or correlative light and X-ray tomography [6].

Molecular localisation in EM has been possible for many years [7] although often through difficult techniques, for example the use of antibodies coupled to different sized gold particles in ultra-thin sections. However, in general this involves chemical fixation and the incubation of antibodies on 60–100 nm ultra-thin sections, which provides a very limited 2D view of the specimen and has very poor detection efficiency. Extending this method to three dimensions is even more challenging, requiring multiple sections to be labelled, imaged and then registered into a single volume.

Correlative fluorescence and ultrastructural microscopy have successfully been applied to the localisation of molecules in plastic sections [8–10]. Using such sections, it has even been possible to carry out super-resolution fluorescence imaging using Single Molecule Localisation Microscopy (SMLM) techniques [11–13]. However, to date such approaches have all involved chemical fixation, which is subject to potential artefacts.

An alternative approach, rapid cryo-freezing, does not suffer from the same potential morphological artefacts. Compared to chemical fixation, cryo-fixation offers far superior preservation of local chemistry and delicate cellular structures. Therefore, cryo-fixation has now become the gold standard for preserving structures or even snapshots of rapid dynamic processes within cells [14, 15]. Cryo-preservation can also enable the determination of molecular structures within whole cells [16, 17]. It has the additional advantage of enhanced dye photo-stability at low temperatures [18], leading to lower photo-bleaching rates. This permits the accumulation of images with high signal-to-noise ratio, as imaging can be maintained over a large number of frames.

A major challenge in correlative imaging at cryogenic temperatures is how to implement high resolution fluorescence microscopy. The most advanced commercial cryo fluorescence microscopes allow imaging to a resolution of approximately 400 nm using low numerical aperture objectives. Such resolution is insufficient to relate the distribution of molecules to the ultrastructure on scales of 5–50 nm. This means there is a gap in volumetric resolution of several orders of magnitude, which severely limits the usefulness of the correlative approach. A few cryogenic super-resolution demonstrations, to date, have been performed on specialised microscopes with limited accessibility [12, 18].

We present the cryoSIM system as a solution to these technical challenges. This microscope is situated at the Diamond Light Source correlative cryo-imaging beamline B24 in the UK [19]. CryoSIM combines super-resolution structured illumination fluorescence microscopy at cryogenic temperatures with correlative ultrastructural imaging. The technique enables 3D imaging of volumetric specimens at resolutions beyond those achievable with the best confocal imaging, making it an excellent complement to Soft X-ray Tomography (SXT) [20] or EM at cryogenic temperatures. The cryoSIM produces 3D images of volumes over 10 *µ*m in thickness, which is well matched to the penetration depth of the SXT, and can detect fluorescent molecules with wavelength dependant resolutions of up to 200 nm, providing close correlation to the imaged ultrastructure. The design of the cryoSIM instrument allows careful control of sample temperature to maintain sample preservation in vitreous ice by inhibiting formation of ice crystals throughout the process. After fluorescence imaging, the same samples are kept preserved in liquid nitrogen and transferred for ultrastructural studies using SXT (established correlative scheme) or EM (correlative scheme under development). This correlated imaging platform can be applied to many biological samples to gain insights into structure and function in an effort to elucidate molecular mechanisms in biomedical and life science research.

## 2. Microscope design

### 2.1. Choice of super-resolution method

Structured illumination microscopy [21, 22] was chosen as the mechanism for obtaining super-resolution in the cryoSIM system. 3D-SIM offers several advantages over other fluorescence super-resolution techniques for volumetric imaging of thick samples. It has very good out-of-focus light suppression, resulting in high contrast images even in samples with thicknesses of 10 *µ*m or more. 3D-SIM requires relatively low illumination intensities compared to Single Molecule Localisation Microscopy (SMLM [11, 23, 24]) or STimulated Emission Depletion microscopy (STED, [25, 26]). Therefore, 3D-SIM offers reduced sample heating and lowers the risks of ice crystal formation and sample damage. While 3D-SIM requires a large number of raw images to generate the reconstructed super-resolution image, this number is far smaller than that required for SMLM, and there is no requirement for high power densities to saturate the stimulated emission (as in STED) or blinking (as in some forms of SMLM) [27]. Finally, and crucially, 3D-SIM is readily applied to multi-channel imaging using a range of common dyes or fluorescent proteins. Therefore, 3D-SIM provides a flexible mode of super-resolution microscopy that can be readily applied to a very wide range of cryo preserved biological specimens and problems.

### 2.2. Cryogenic specimen stage

In order to image fluorescence samples at cryogenic temperatures, we built a custom optical microscope around a commercial cryostage (Linkam CMS196M LED Cryo Correlative Stage). The entire system was designed to maintain good temperature stability, enabling precision imaging, maximising mechanical stability, minimising thermal drift and maintaining the essential vitreous state of the specimen. Image drift is cyclical and almost entirely in the X direction at 2 nm/s (measured using time-lapse of fluorescent beads at cryo temperatures; see supplemental Fig. S1 and Fig. S2). It is essential to prevent ice crystals forming in the specimen, as ice crystals can disrupt the sample’s structure, and introduce artefacts that can affect light-, electron-, and X-ray imaging. To achieve this goal, we used a rapid plunge-freezing protocol that locks the water in the sample into a vitreous state, and then kept the sample below the glass transition temperature of its aqueous component: approximately 130 K (−140°C). It should be noted that freezing thick samples is challenging and approaches such as high pressure freezing might be required.

The commercial cryostage was designed for use in microscopy and ensures the sample is always maintained in contact with a cold block immersed in liquid nitrogen, Fig. 1. The microscope uses a long working-distance air objective (Nikon, TU Plan Apo BD 100× 0.9 NA, 2 mm working distance, so that the air gap between the sample and relatively warm objective provided effective thermal insulation and prevents the objective from heating the specimen. We also control the intensity of the illumination from the fluorescence excitation laser light sources, as light absorption can cause significant local heating and can result in ice formation or even thermal damage.

**Fig. 1.**
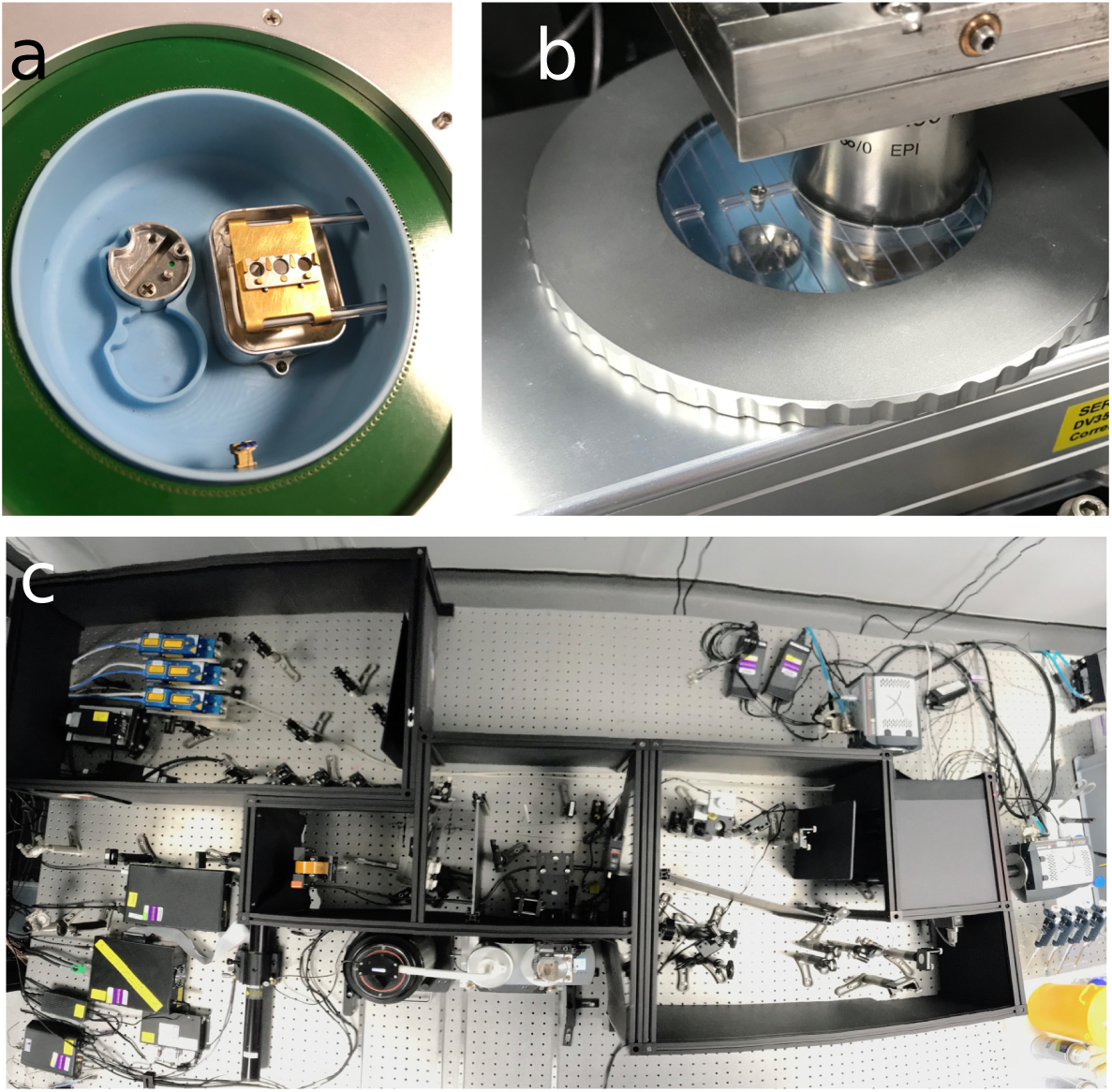
CryoSIM sample stage: (**a**) Closeup view of the cryo sample stage, with the 3 position grid holder on top of the copper bridge, which dips into the liquid nitrogen bath, (**b**) A view of the objective lowered into the cryostage in its imaging position with the cover in place to reduce sample warming, and (**c**) a view of the system as a whole from above with the cryostage central at the bottom. The optics are usually enclosed in the black boxes shown here with their lids removed.

### 2.3. Optical system design

A schematic of the cryoSIM microscope is shown in Fig. 2. This includes the optical elements for producing the structured illumination and the emission path for imaging the fluorescence.

**Fig. 2.**
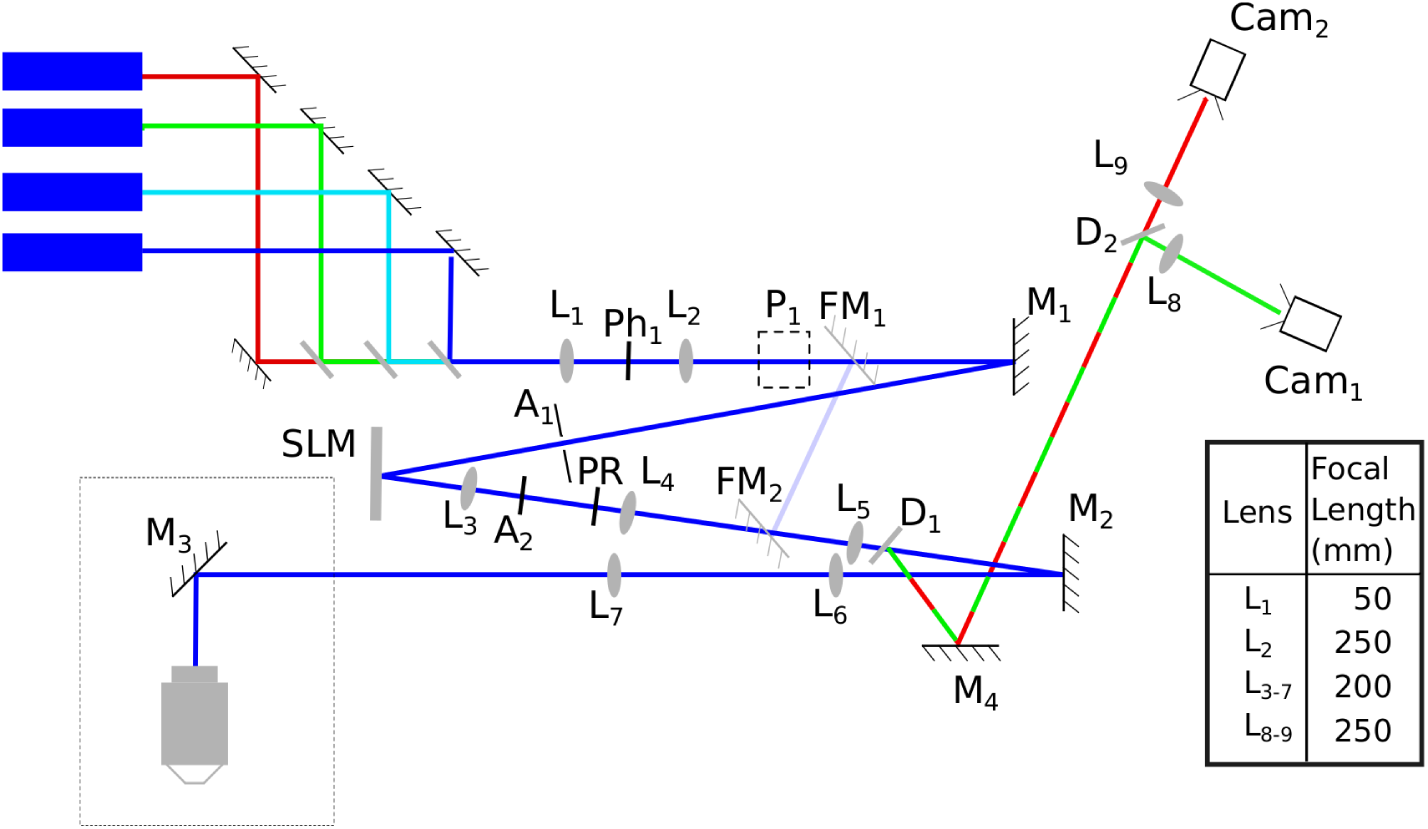
A schematic of the cryoSIM microscope, with lenses (L), apertures (A), mirrors (M), pinholes (Ph), dichroics (D), periscope (P), polarisation rotator (PR), and cameras (Cam) shown. The four lasers have wavelengths of 405, 488 and 647 nm (Omicron DeepStar lasers) and 561 nm (Cobolt Sapphire laser) were combined and passed through the telescope L_1_ and L_2_ with a pinhole (Ph_1_) at its focus to clean up the beam profiles. The beam was reflected reflected towards the SLM by M_1_, through an aperture A_1_. Light reflected from the SLM was refocused by L_3_ to aperture A_2_ which removed high diffraction orders generated by the SLM. The polarisation was rotated by the PR and the telescope formed by L_4_ and L_5_ reimaged the diffracted spots onto M_2_, after L_5_ the excitation light passed through the dichroic D_1_. The beam continued through telescope L_6_ and L_7_ via M_3_ to the back-pupil of a 100× 0.9 NA air objective and focused into the sample. The fluorescence emission was collected by the objective and reflected from M_3_, passed back through L_7_ and L_6_, was reflected M_2_ to dichroic D_1_, where the fluorescence emission was reflected off M_4_ to the dichroic D_2_. This dichroic split the emitted light between two cameras, Cam_1_ and Cam_2_, via imaging lenses L_8_ and L_9_. The dotted box in the bottom left hand side was at 90°to the plane of the rest of the diagram, so while most of the optics are in the horizontal plane, the objective is vertical and pointing downwards. There are two flip mirrors FM_1_ and FM_2_ which bypass the SLM to allow widefield illumination. Lens focal lengths are shown in the table.

Lens pairs are setup as 4 *f* telescopes, except for L_8_ and L_9_ which are further down the optical path due to physical space constraints. However, the beam is infinity focused at this point so this design compromise has minimal effect on the image quality. Most of the optical design choices are standard options that would be used in any optical system, for example all lenses were chosen as achromatic doublets. However, there are some specific design optimisations covered below that significantly improve the imaging using structured illumination.

Lens orientation was chosen to optimise image quality. As a general rule, in order to minimise aberrations, lens orientation should be chosen to minimise the deviation of the light-lens interaction angle from the normal. However, in this microscope, the excitation light and the emitted fluorescence are in reciprocal focal states: when the excitation light is infinity focused between two lenses in a telescope, the emitted light will be coming to a focus and then spreading out again. Sections of the optical system that contain either only excitation or only emission paths can be optimised with a particular lens orientation. Lenses which are in parts of the system that include both paths, i.e. those between the dichroic and the objective, were optimised for the emitted light to minimise image aberrations.

### 2.4. Spatial light modulator and pattern optimisation

Effective 3D-SIM operation requires illumination patterns with high contrast structures at the limit of resolution. In order to generate the illumination patterns, we used a nematic liquid crystal on silicon Spatial Light Modulator (SLM, Meadowlark optics, 512×512 SLM Model PDM512), which enabled grayscale phase modulation. The SLM allows a large fraction of the light to be transferred into the first order diffraction spots even in stripes with a width close to 2 times the pixel size, unlike a binary ferroelectric SLM, which exhibits lower efficiency. Polarisation control is also vitally important for optimal pattern contrast, as near-perfect linear polarisation is required in the focus. For this reason, between the SLM and sample, silver-coated mirrors (Thorlabs PF10-03-P01) were chosen in preference to broadband dielectric mirrors. This results in better preservation of linear polarisation, optimising pattern contrast at the expense of a marginal decrease in reflection efficiency. At other points in the system, where depolarisation was not important, broadband dielectric mirrors (Thorlabs BB1-E02) were used to maximise reflection efficiency. The multi band dichroic (Chroma ZT405-488-561-647-22.5deg) was a custom design that operated at an incidence angle of 22.5°rather than the more conventional 45°. The lower incidence angle produced a sharper cut-off between reflection and transmission, so that the laser transmission bands could be narrower and a greater proportion of the fluorescence emission could be detected. Additionally, this dichroic was used in an orientation that was inverted, compared to those on a conventional fluorescence microscope, in that the illumination was transmitted and the emission was reflected. The reflection mode is slightly more efficient, leading to an improved detection efficiency.

Moreover, in transmission, the dichroic has less effect on the light polarisation in the illumination path, thus improving pattern contrast.

To strengthen the information content in the highest lateral frequencies we introduced on the SLM a sinusoidal phase pattern with a base offset that was empirically chosen to generate relative beam intensities of 1 in the central beam and 1.3 in the +1^*st*^ and −1^*st*^ order diffracted beams as previously suggested [21]. The SLM can be used to adjust the pitch of the sinusoidal pattern to account for the variation of the diffraction limit with wavelength. We also included a polarisation rotator (Meadowlark LPR-100-*λ*) in the optical path, to ensure that the polarisation was set for each pattern orientation to maintain maximum contrast. This required the polarisation to be linear and orientated radially in the back focal plane of the objective. The polarisation rotator was calibrated for each wavelength and stripe orientation in order to provide the correct radial polarisation.

To ensure that the illumination patterns were in a focal plane coincident with the camera image plane, we placed the first lens after the SLM on a linear stage, whose direction of motion was aligned with the optical axis. This allowed adjustment of the focus of this lens without realignment of the rest of the optical setup. We found that shifting the illumination pattern at the sample in the Z direction by one cycle required about 15 mm of linear translation of this lens (Fig. 2, L_3_)

### 2.5. Camera configuration

Two cameras (Andor iXon Ultra 897) were employed to acquire different channels in sequence, thus allowing fast image acquisition. A second channel could be collected while the image of the first channel was read out from the camera. Such a fast imaging regime helped ensure that the limited residual thermal drift had minimal impact in the images produced. As each channel was captured by a separate camera behind a filter wheel (Thorlabs FW102c), the system required careful alignment of the camera positions so that they were parfocal. The optical design also allowed conventional widefield fluorescence imaging using one flip mirror (Fig. 2 FM_1_, Newport 8892 Flipper Mount) to bypass the SLM and polarisation optics, and another (Fig. 2 FM_2_) to couple back into the main beam path.

### 2.6. Hardware Synchronisation

During experimental image collection the system was coordinated via hardware triggering from a Digital Signal Processor (DSP, Innovative Integration M67-160 with an A4D4 daughter board) [28]. This hardware triggering approach, adapted from Carlton et al. and their OMX microscope, is able to produce both digital triggers and analogue signals with precise timing. The control software produced a timing table used by the DSP to digitally trigger the lasers, cameras, and the SLM, which had preloaded patterns. The DSP also produced specific analogue voltages for the piezo Z stage and the polarisation rotator at specific times. The timing table was carefully constructed to allow the correct timing intervals for devices such as the piezo, polarisation rotator, and SLM which take significant time to reach the correct state before the next image acquisition occurs. Code to run this hardware, both the Python code and the binary object that runs on the DSP, is available online [29]. However, the boards are difficult to source and we are currently working on a replacement solution.

### 2.7. Image acquisition

Image stacks were acquired as either single colour images, or two colour images. Two colour images interleaved the colour channels, so images in each channel were taken one after the other. SIM stacks were acquired by taking 5 phase steps at one angle then changing the angle and taking a further 5 phase steps. After 3 angles were taken in one or two colours the focus was stepped to the next Z position and this was repeated. There are then 15 images per Z plane, per colour.

### 2.8. Objective mount and stage Z-control

To improve ease of use and workflow during imaging, we have designed custom parts to hold the objective and to provide coarse movement of the cryostage along the optical axis. A piezo stage enables us to precisely position the cryostage in Z, but only offers a 200 *µ*m range of motion. We found that this was inadequate for convenient use, due to variation in sample holder thicknesses and because grids can demonstrate significant curvature, requiring larger excursions to bring the region of interest coincident with the focal plane. To enable larger shifts, the combined piezo and cryostage assembly is mounted on a platform that is driven by a leadscrew, turned by a hand-operated wheel. The wheel is indexed, with steps between adjacent indices moving the sample by 50 *µ*m, and giving a total range of a few millimetres.

For imaging, the objective must be inserted around 15 mm into the sample chamber to bring its focal plane into the sample plane. The objective is mounted in a plate which forms the upper half of a separable kinematic mount, and which can be raised and lowered via a cam attached to a lever arm. When the objective is raised, the entire cryostage assembly can be slid forwards on linear bearings, towards the user and away from the imaging position. The user is then able to undertake sample transfers without hindrance, and with reduced risk of knocking components in the optical train out of alignment. After a sample is loaded, the cryostage is slid back into the imaging position where it is held in place by a detent. The cam lever is then used to lower the objective plate so that hardened steel balls bonded to the plate sit in V-grooves on a lower plate, ensuring reproducible and stable positioning. The lower plate sits between two sets of fine thread screws and stiff leaf springs, so that it may be translated within the assembly along two axes lying perpendicular to the objective axis. This allows precise lateral positioning of the objective to bring it into alignment with the rest of the optical path. Technical drawings for these custom assemblies are listed in the supplement 1 Table S1 along with the pdfs, and the CAD files are available upon request.

### 2.9. Software and control

A major objective of the cryoSIM design was to enable ease of use for experienced microscope users and to allow others to adopt the design, modify it, and develop new bespoke workflows for specific applications. With this in mind, the system was assembled almost entirely from commercially available parts and using standard optical components. This aim extends to the control software and so we have chosen to use Python-Microscope and Microscope-Cockpit, both of which are free and open source software. Microscope-Cockpit provides, an easy to use, user friendly interface to complex automated microscopes [30] and is based on software from UCSF [28]. It is built on top of Microscope, a Python library for the control of individual microscope devices [31]. While developing cryoSIM, we have made extensive modifications to both Microscope-Cockpit and Python-Microscope. These modification have been merged into their code base and are included on their releases. Finally, synchronisation of the individual devices during an experiment is performed with TTL signals and analogue voltages via a DSP board controlled by Microscope-Cockpit.

## 3. Materials and Methods

### 3.1. Bead sample preparation

All samples were imaged on carbon coated 3 mm gold EM finder grids (Quantifoil). Bead test and calibration samples were made by adding 2 *µ*l of 175 nm fluorescent bead solution(Thermo Fisher PS-Speck), diluted by a factor of 10^5^ in distilled water and added to Quantifoil grids. These samples were allowed to air dry. They were then cooled on the Linkam cryostage.

### 3.2. Cellular sample preparation

Cellular images were taken of HeLa cells grown on carbon coated gold EM grids under standard tissue culture conditions. Initial images were collected from cells labelled with MitoTracker Red FM (Thermo Fisher) at 100 nM and LysoTracker Green DND-26 (Thermo Fisher) at 50 nM after 30 minutes of incubation, to give green lysosomes and red mitochondria. The grids with live cells on them were then blotted and plunge frozen in liquid nitrogen cooled liquid ethane (Leica EM GP2) before being transferred to liquid nitrogen storage. Once frozen, grids were preserved in liquid nitrogen or at cryogenic temperatures on the imaging systems to prevent thawing and detrimental ice crystal formation. Imaging shows ice layers are less than 0.75 *µ*m in thickness (see Supplemental Fig. S3).

*Drosophila melanogaster* primary post-embryonic haemocytes (plasmatocytes) were extracted from Drosophila 3^*rd*^ instar larvae and grown in Schneider’s Drosophila Medium (Thermo Fisher) on 200 mesh carbon coated gold TEM F1finder grids (Quantifoil). CryoSIM data were collected from cells labelled with ERtracker Green (Thermo Fisher) at 500 nM, LysoTracker Red DND-99 (Thermo Fisher) at 50 nM and MitoTracker Red FM at 250 nm added in the growth medium for 45 minutes. Following incubation, samples were blotted with filter paper (0.5 s) to remove the excess liquid, and plunge-frozen in liquid ethane (Leica EM GP2).

Further samples were prepared from stable and inducible HeLa cells expressing mov10-YFP [32] to label p-bodies. HeLa cells were induced to produce mov10-YFP by supplementation of the medium with 1 *µ*g/ml doxycycline and then infected with sindbis virus (SINV-mCherry, see below [33]) and then incubated with MitoTracker Deep Red (Thermo Fisher) at 125 nM for 30 minutes before being plunge frozen as above for imaging.

We used the SINV clone pT7-SVmCherry to generate the SINV-mCherry suspension. The plasmid pT7-SVmCherry was generated by inserting mCherry after the duplicated subgenomic promoter in pT7-SVwt [34]. To obtain SINV-mCherry viruses, the pT7-SVmCherry plasmid was first linearised with XhoI and used as a template for in vitro RNA transcription with HiScribe T7 ARCA mRNA kit (New England Biolabs, #E2065S). Transcribed genomic RNA was transfected into BHK-21 using Lipofectamine 2000 reagent (Invitrogen, #11668027). Viruses were collected from the supernatant 24 hours later and cleared by centrifugation at 2000 rpm for 3 minutes followed by filtration with 0.45 *µ*m PVDF syringe filter units (Merck, #SLHV033RS). Cleared supernatants were titrated by plaque assay using BHK-21 cells.

### 3.3. SIM image calibration

Calibration Optical Transfer Functions (OTFs) were created by imaging single isolated 175 nm fluorescent beads (Thermo Fisher PS-Speck), with no other bead within 30 *µ*m. These single colour beads were imaged by a standard SIM image protocol of 5 phases and 3 angles of stripes with 0.125 *µ*m Z spacing and a stack height of 8 *µ*m. Images were then cropped to 256×256 pixels around the bead centre, and a single angle of stripes used for processing into OTFs using SoftWoRx. OTFs for all channels were created separately using beads with the optimal fluorescent emission.

### 3.4. SIM image collection and reconstruction

For super-resolution acquisition, images were taken sequentially with 5 different phase shifts of the sinusoidal pattern, at 3 orientations, repeated for every Z position in a full 3D stack. After acquisition the 15 resulting images per Z plane were then computationally recombined to produce an image with up to double resolution in all 3 dimensions, as has been previously described for 3D-SIM [21, 22, 35, 36]). Reconstructions were performed in SoftWoRx (GE) and multi-channel images were aligned using 4-colour fluorescent bead (Thermo Fisher TetraSpeck) reference images, and Chromagnon [37] to calculate and apply the relevant transforms. The raw and reconstructed data were analysed with the open source ImageJ plugin SIMcheck [38, 39] to estimate the resolution enhancement, ensure the results were realistic, and contained minimal artefacts. Widefield images were synthesised by averaging all 15 images from a single Z plane into a single image, fully sampling the image plane, and producing widefield images without having to collect more data, while also avoiding any alignment issues when comparing widefield and SIM images (e.g., see Fig. 4). SIMcheck was used to generate radial averaged power spectral density plots and image decorrelation analysis [40], implemented in ImageJ using default parameters, was used to numerically assess resolution in cellular context.

**Fig. 3.**
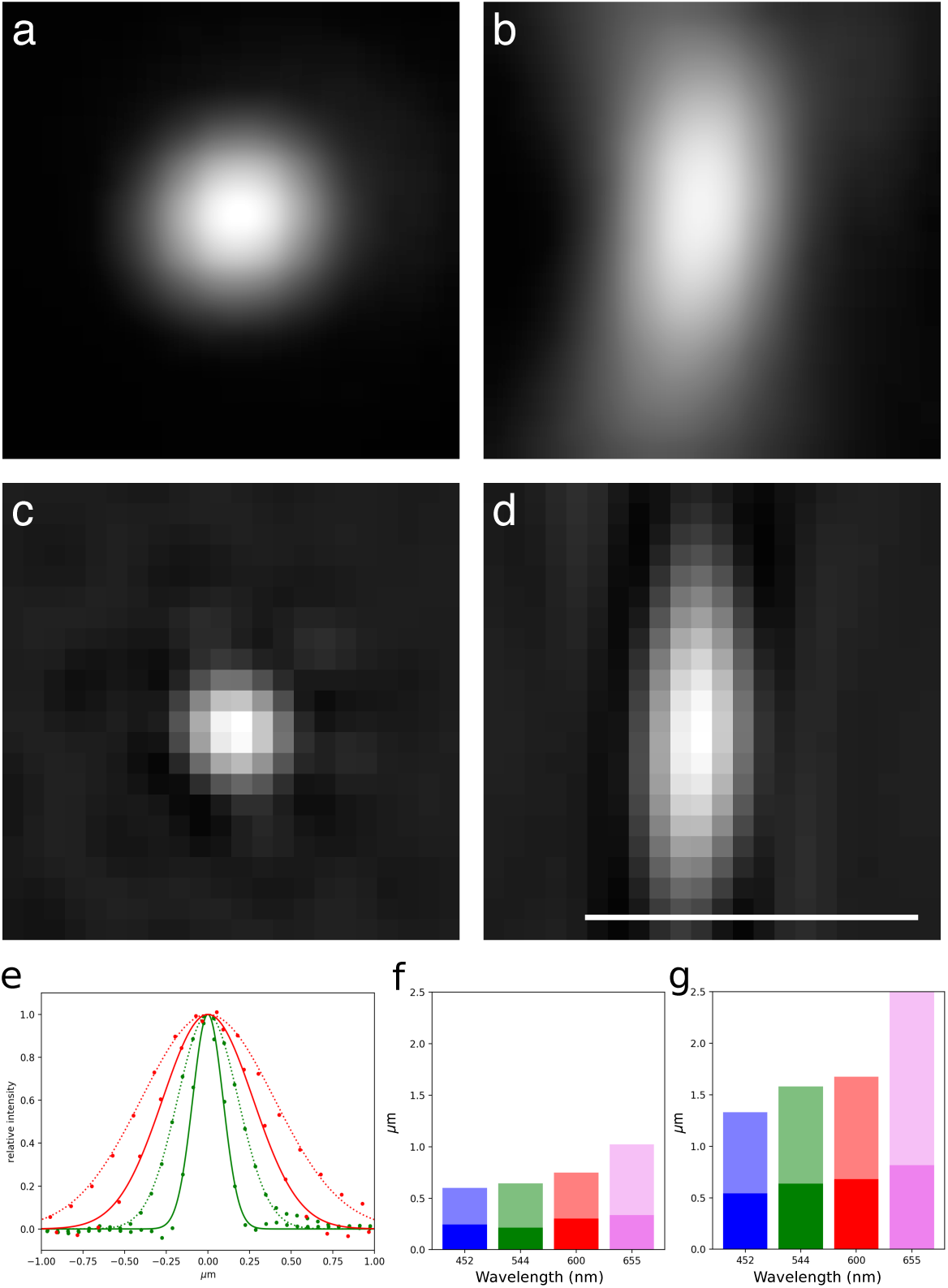
Widefield and SIM point spread functions: Images of a single 175 nm diameter fluorescent bead in focus (**a, b**) widefield in XY and XZ planes and (**c, d**) SIM in XY and XZ planes respectively. The beads were imaged with 488 nm excitation and 544/24 emission filter. (**e**) line scans through this bead with measured values as points and Gaussian fits as lines. The point spread function in the lateral (XY) direction is in green and the axial (Z) in red. (**f, g**) measured FWHM of the Gaussian fits in the lateral (**f**) and the axial (**g**) directions with SIM data as dark bars and widefield data as fainter bars. Scale Bar 1 *µ*m.

**Fig. 4.**
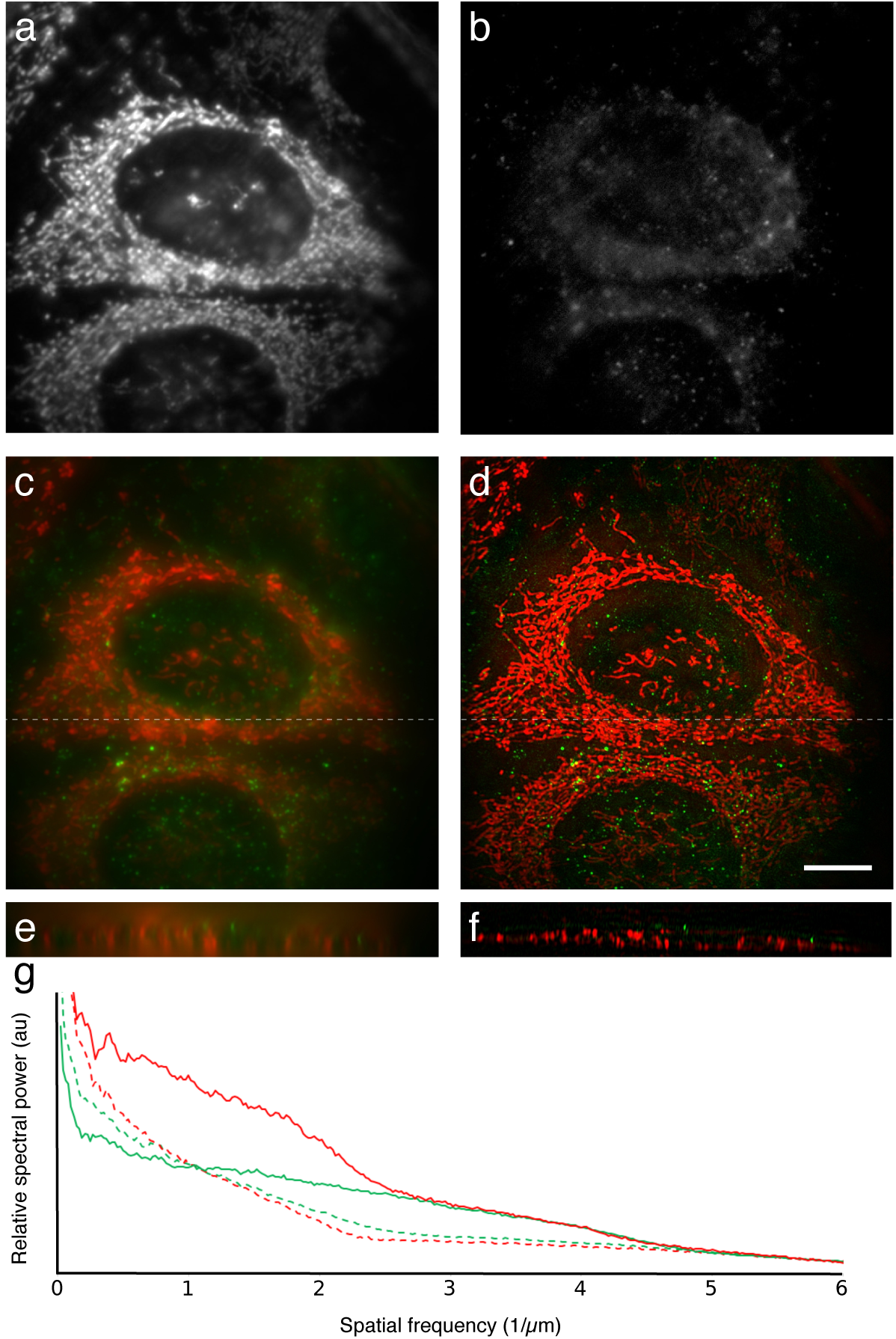
CryoSIM images of HeLa Cells stained with Mitotracker red and Lysotracker green excited at 488 nm and 561 nm respectively. (**a**) Raw Mitotracker red signal, (**b**) raw Lysotracker green signal, (**c, d**) Widefield and SIM reconstruction maximum intensity projection over 2.75 *µ*m depth, (**e, f**) XZ slice of Widefield and SIM reconstructed stack, at the position marked by the dashed lines in **c** and **d**, to show the increased Z resolution. (**g**) Spectral power density plot to show the increase in information content between widefield (dashed lines) and SIM reconstructions (solid lines), especially in the range 2.5–5 (0.4–0.2 *µ*m). Image decorrelation analysis gives widefield resolutions as 508 nm and 606 nm and SIM resolutions as 216 nm and 345 nm in green and red respectively. Scale bar 10 *µ*m.

X-ray tomograms were aligned and reconstructed using IMOD [41] and the same processing as in Kounatidis et al. [20].

## 4. Results

### 4.1. Bead images demonstrate resolution at cryo temperatures

The microscope produced conventional images near the theoretical diffraction limit. Fig. 3 shows images of a 175 nm fluorescent bead using blue excitation at 488 nm and a green emission wavelength centred on 544 nm. The Full Width Half Maximum (FWHM) dimensions of these images were 430 nm in the lateral direction and 940 nm in the axial direction.

The super-resolving abilities were tested in 3D-SIM mode. The illumination patterns had pitch close to the resolution limit: 330 nm for 405 nm excitation, 396 nm for 488 nm excitation, 451 nm for 561 nm excitation, and 520 nm for 647 nm excitation. Reconstructed bead images in the green, for comparison to the widefield data above, showed FWHM of 210 nm in the lateral direction and 640 nm in the axial (Fig. 3e-g).

These observations show conclusively that by combining super-resolution imaging with cryo-preserved samples, we are able to image outstandingly well preserved structures at resolutions beyond those achievable by conventional fluorescence imaging in fixed or live cells.

### 4.2. High-fidelity cellular imaging in example samples

In order to demonstrate the biological application of cryoSIM we imaged HeLa cells which were fluorescently labelled in the mitochondria and lysosomes while live and then plunge frozen. HeLa cells were grown using standard tissue culture conditions on carbon film coated gold EM grids. The cells were labelled by adding Mitotracker red and Lysotracker green (see materials and methods for details). Mitotracker is a fluorescent dye that specifically binds to mitochondria while Lysotracker binds to lysosomes, both work well in living cells.

Grids were transferred from cryo storage to the cryoSIM grid holder on the Linkam stage kept at cryogenic temperatures. Brightfield transmission imaging was carried out to locate individual cells with good overall morphological preservation and no obvious grid surface or cell sample disruption. The Microscope-Cockpit program enables large regions or even the entire grid to be imaged through the use of a spiral mosaic pattern, without the need to change objectives. Cells were marked within the brightfield global view panel, for convenient revisiting for 3D-SIM imaging with fluorescence illumination. 3D-SIM data was collected on a number of representative cells across each grid and processed to produce super-resolution reconstructions (Fig. 4). The results indicate that both contrast and resolution are substantially improved in the SIM reconstructions compared to the widefield images. The plot of spectral power density (Fig. 4g) shows the clear gain in information content above the classical resolution limit and decorrelation analysis ([40]) shows resolutions increasing from 508 nm to 216 nm in green and 606 nm to 345 nm in the red, demonstrating conclusively that 3D-SIM imaging is achieved at liquid nitrogen temperatures.

Additionally, to show the versatility of cryoSIM across biological cell types and ability to image a wide range of dyes in 3D super-resolution, we imaged *Drosophila melanogaster* primary post-embryonic haemocytes in 3 colours. Live cells were stained for the ER in green, lysosomes in red and mitochondria in farred. These triple labelled cells were then imaged and SIM reconstructions are presented in Fig. 5.

**Fig. 5.**
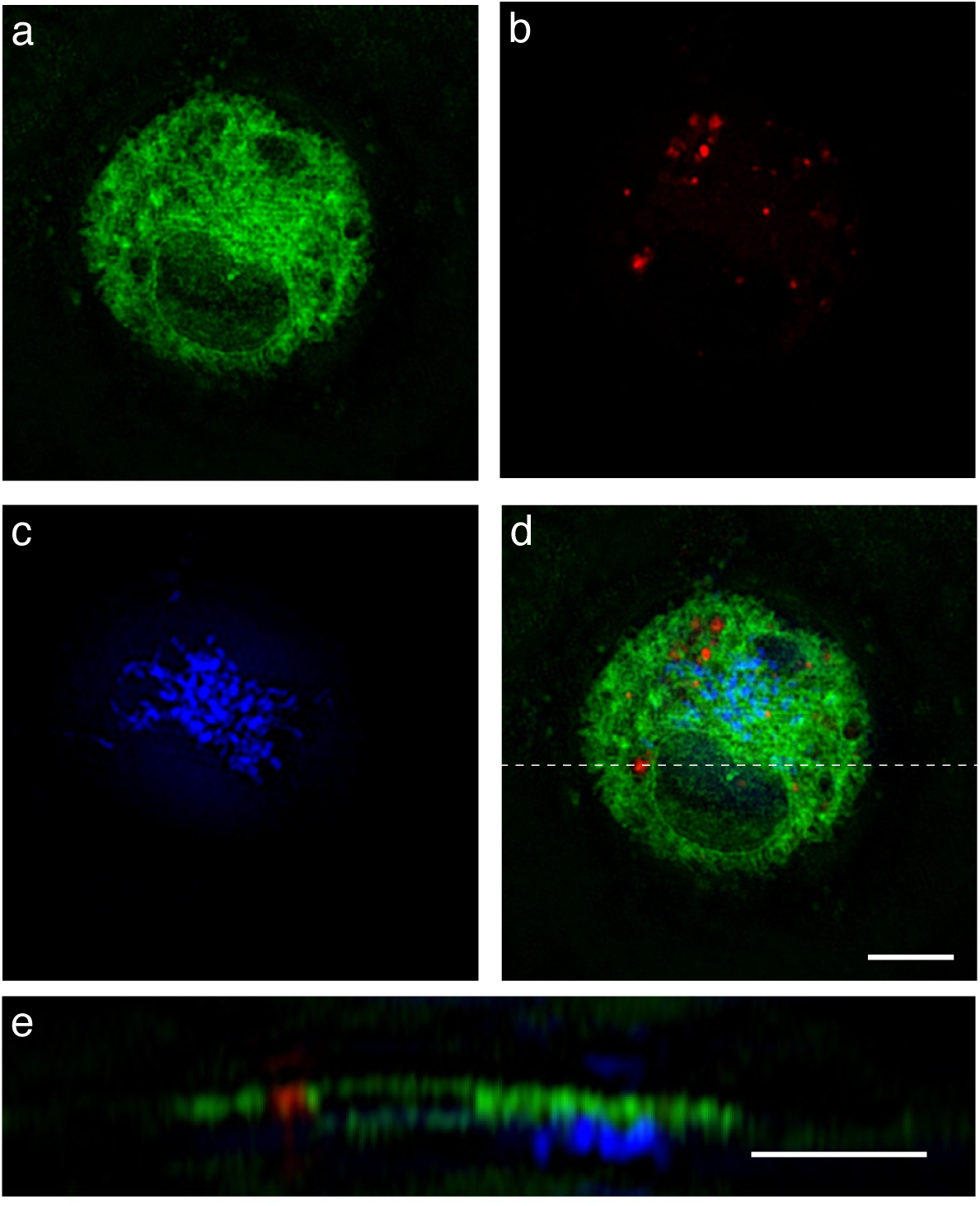
CryoSIM three colour imaging of ER, Lysosomes and mitochondria in *Drosophila melanogaster*. (**a**) ER labelled in green with 488 nm excitation, (**b**) Lysosomes labelled in red with 561 nm excitation and (**c**) Mitochondria labelled in the farred with 647 nm excitation (shown in blue), (**d**) a merged image. All images are maximum intensity projections over 750 nm depth. (**e**) XZ projection at the position marked by the dashed line in **d**, note this is at twice the scale of the other panels. Image decorrelation analysis produced lateral resolutions of 217 nm, 248 nm and 307 nm in the green, red, and farred respectively. Scale bars 5 *µ*m.

### 4.3. Correlative imaging using cryoSIM and soft X-ray microscopy and tomography

To facilitate a convenient and generally applicable workflow for correlative imaging of whole cells, we built the cryoSIM microscope at the soft X-ray tomography (SXT) beamline at Diamond Light Source. Although SXT typically offers a lower resolution than electron microscopy, it has the advantage that that it delivers up to 25 nm imaging while its soft X-rays can penetrate ice, or vitreously frozen biological samples, far deeper than electrons: thick samples can be imaged as a tilt series, without resorting to laborious physical sectioning techniques. The 3D structure of the sample is then determined using standard computational tomographic reconstruction algorithms [42]. Correlative imaging has given insights into a range of cellular processes, and the ability of both cryoSIM and SXT to handle the same thick sample opens up the application of this type correlative cryo-imaging to the study of both intra- and inter-cellular processes in larger tissues, as well as isolated cells.

We imaged stable and inducible HeLa cells expressing MOV10-YFP and infected with sindbis virus on both the cryoSIM and the cryoSXT instrument to demonstrate the correlative imaging potential of our platform. First cells were imaged in fluorescence on the cryoSIM setup and then large areas of the grid were imaged as a simple 2D X-ray mosaic. These images could then be aligned to ensure we were imaging the same region of the same cell (Fig. 6). Once a suitable location was found, X-ray tilt series were collected. The alignment information from the 2D X-ray image allows correlation of the 3D image stacks between cryoSIM and X-ray tomography (Fig. 7). The clear imaging of the same region, at different resolutions is shown by the comparisons between the tomogram and the fluorescence in XY images (Fig. 7f-h). The YZ sections, chosen as the tomogram is gathered around the Y axis so the resolution is optimised in this direction, show the almost isotropic resolution in the tomogram section (Fig. 7i) but much more elongation in the Z direction in the fluorescence images (Fig. 7j-l).

**Fig. 6.**
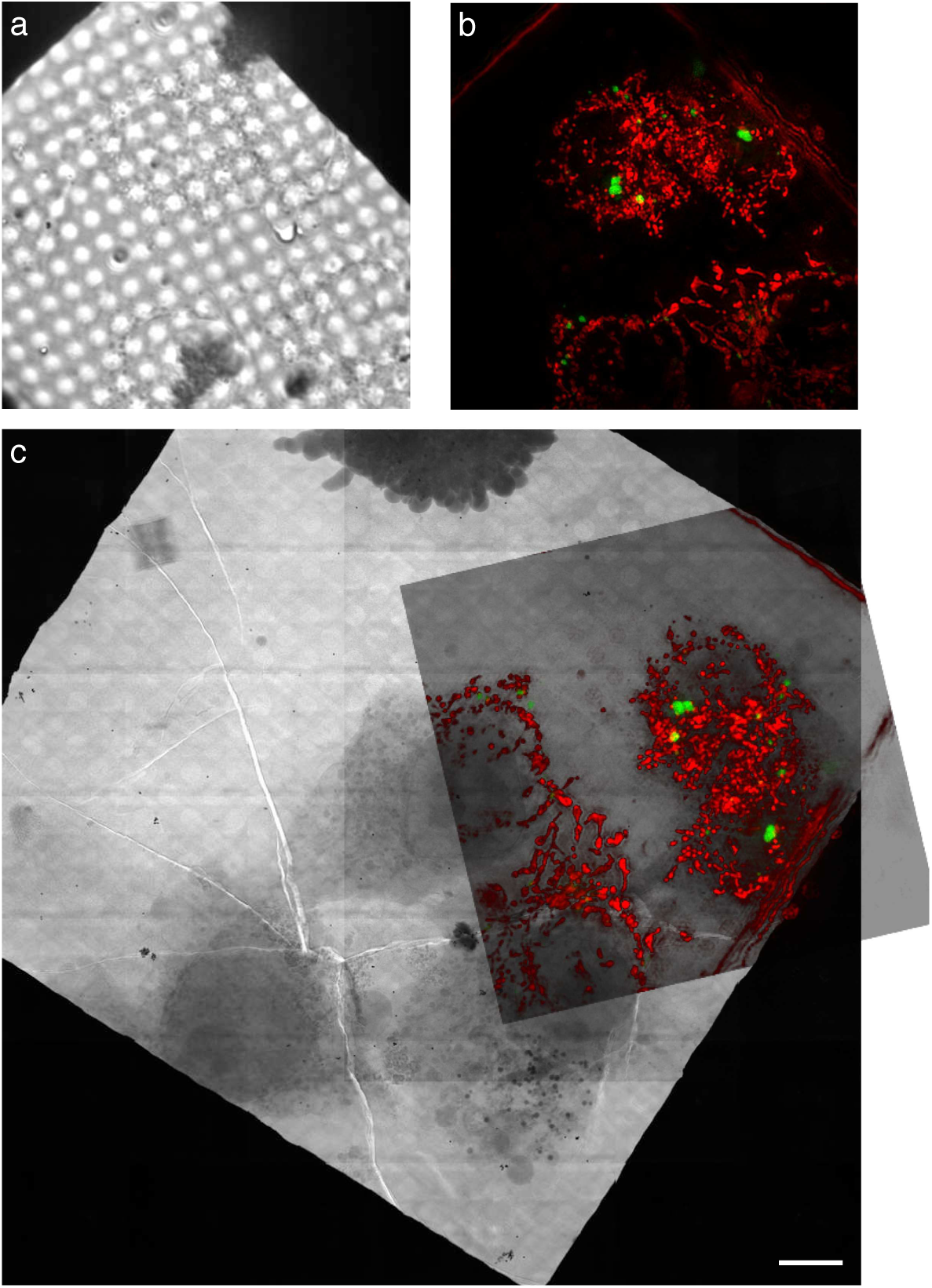
Correlated cryoSIM and X-ray microscope images. HeLa cells expressing mov10-YFP and labelled with MitoTracker Deep Red. (**a**) transmission image from cryoSIM microscope, (**b**) SIM reconstruction fluorescence image from the same region. A maximum intensity projection over 3.125 *µ*m with mov10-YFP in green and MitoTracker Deep Red in red, XZ projections of this data is shown in supplemental Fig. S4. (**c**) a mosaic from the X-ray microscope with the semi-transparent SIM fluorescence reconstruction rotated and scaled to its correct location, clearly showing the same set of cells. Scale bar 10 *µ*m.

**Fig. 7.**
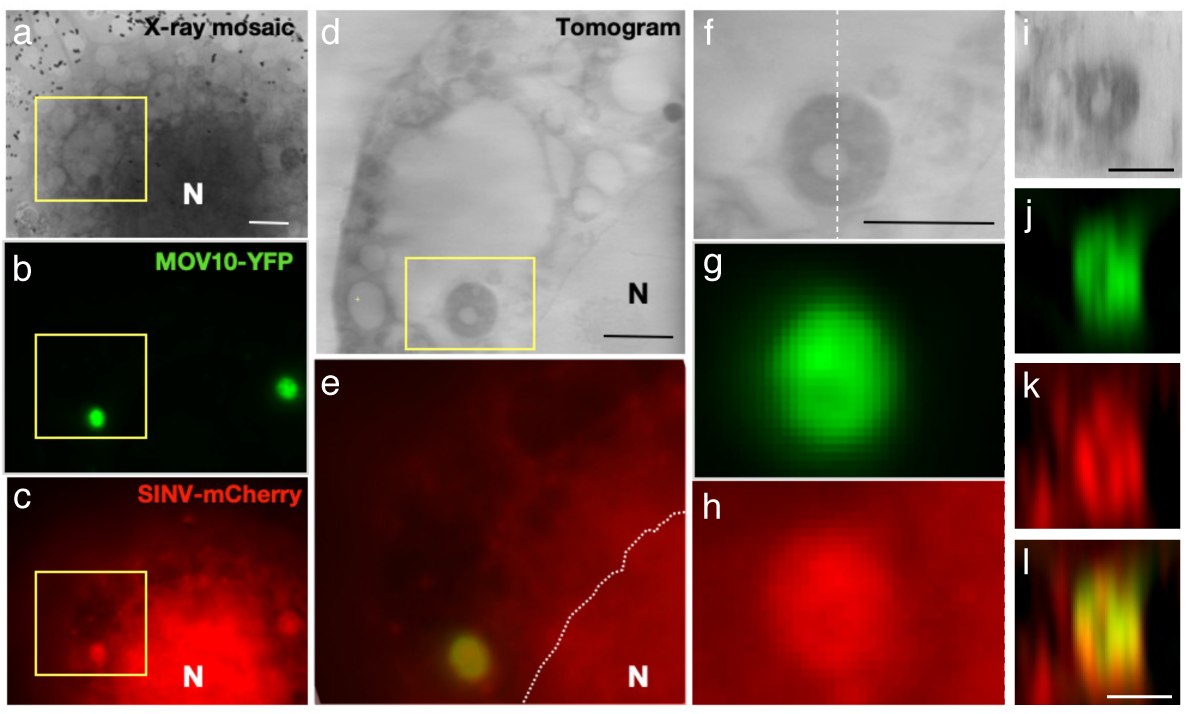
Correlated cryoSIM and X-ray tomogram. HeLa cells expressing mov10-YFP and infected with SINV-mCherry. (**a-c**) A section of the overview X-ray mosaic image (**a**) with maximum intensity projections of SIM reconstructions of the same area for mov10-YFP (**b**), and SINV-mCherry (**c**). (**d, e**), Magnified images of the boxed region from **a-c** in a slice of the X-ray tomogram (**d**) and merged channels from the SIM reconstructions (**e**). (**f-h**) further magnifications of the boxed region in **d**, clearly showing a dense structure (**f**) which colocalises with both mov10 staining in green (**g**) and SINV-mCherry in red (**h**).(**i-l**) YZ sections, along the line shown in **f**, of the pbody region in the x-ray tomogram (**i**), mov10 in green (**j** and **l**) and SINV-mCherry in red (**k** and **l**). The nucleus is marked as N with its boundary indicated with a dotted line in panel **e**. (Scale bars **a, d** = 5 *µ*m; **f, i** and **l** = 2 *µ*m)

## 5. Discussion

Super-resolution imaging has revolutionised fluorescence microscopy in fixed samples and in many live cell imaging studies. We have developed a super-resolution fluorescence microscope to bring similar benefits to correlative imaging for cryo-preserved samples. The improved resolution substantially increases the information content of the correlative process as it improves the precision of localisation of specific molecules such as proteins and nucleic acids in the ultrastructural morphology.

We chose 3D-SIM as the super-resolution imaging technique of choice for this instrument because it has several distinct advantages for cryo-imaging of general biological samples. First, 3D-SIM requires relatively low illumination intensities, while still giving up to eight times more volumetric information than conventional widefield or confocal fluorescence imaging, a factor of two in each of the three dimensions. In fact, SIM has one of the lowest total light dose of the super-resolution techniques [3], an important factor in cryo imaging, where all light dose must be minimised to prevent sample heating. Although 3D-SIM requires 30 times as many images as widefield fluorescence, 15 images per Z-plane, and the doubled Z resolution requires twice as many Z-planes to satisfy the Nyquist-Shannon sampling theorem, this is relatively low compared to either Single Molecule Localisation Microscopy (SMLM), which requires several thousand images per plane, or STimulated Emission Depletion (STED), which requires GW/cm^2^ depletion laser power [3]. Second, SIM is able to image 3D samples over a relatively large depth, eliminating the need for physical sectioning. Third, imaging can be performed using standard dyes, or fluorescent fusion proteins, and it is easy to image multiple fluorescent channels on a single sample. 3D-SIM is therefore applicable to most cryogenically preserved biological specimens.

Fluorescence imaging at cryo temperatures is usually compromised by limited resolution due to reduced objective NA. By optimising the system to maximise the resolution and light throughput with a long working distance air objective and utilising 3D-SIM, we have been able to image samples beyond the limits of conventional resolution despite the relatively low 0.9 NA objective. The main goal of this setup is to enable super-resolution imaging of fluorescence markers prior to correlative ultrastructural imaging. However there are other significant advantages of imaging in cryo conditions. The low thermal energy dramatically reduces photo-bleaching, giving more than an order of magnitude slower photo-bleaching for the same light exposure. As imaging is almost always photon limited, the reduced photo-bleaching leads to significantly better images than might be expected given the reduced imaging NA available. Rapid cryo freezing of the sample also immobilises fast moving molecules, allowing visualisation of snapshots of rapid processes with no motion blur, or modification of structure during the fixation or imaging process.

However, the optical performance of cryoSIM is significantly worse in cryogenic conditions than at room temperature, due to aberrations produced by the temperature gradients in the gas between the sample and the objective and the lower temperature of the objective itself (supplemental Fig. S5). The design of the microscope includes a mirror in a conjugate plane to the back focal plane of the objective (Fig. 2, M_2_) and we plan to replace this mirror with an adaptive deformable mirror. This mirror will be used to eliminate the aberrations created by cryo-imaging.

Cryo preservation is often used to maximise structural preservation for ultrastructural studies using techniques such as cryoEM and soft X-ray tomography. This instrument is ideally suited to be used in combination with cryoSXT due to both technique’s ability to image thick samples. We have already used the instrument extensively within beamline B24 at the UK synchrotron to correlate cryoSIM and SXT images with a variety of biological samples from a diversity of users. The correlative workflow involved applied to Reovirus studies is published separately in a parallel manuscript [20].

Although we only provide data for correlative fluorescence and SXT imaging it would be relatively easy to extend the current workflow to correlative imaging with electron microscopy. The workflow would be identical until the cryoSIM has been performed and then the sample could either be freeze substituted, and resin embedded similar to refs [8, 12], the sample could be kept in cryo and thin regions like lamella imaged with standard CryoEM techniques [43], or volumes of interest can be located and focused ion beam milling used to create thin lamella to allow tomography [44].

We describe here a first generation instrument, but the microscope has been sufficiently stable and robust to be a fully operational facility instrument at the Diamond Light Source beamline B24. Peer reviewed access has allowed successful data collection for 25 independant imaging projects access with more than 40 days used between February 2019 and March 2020 [20]. The instrument has been used to study a range of biological applications from a number of research groups.

## 6. Conclusions

By developing a convenient super-resolution cryo-microscope platform, we have opened up the investigation of plunge frozen hydrated cells to a higher resolution at a near-native state. These advances will enable many mechanistic molecular insights into biological function. By adding correlative fluorescence workflows specific proteins or nucleic acids can be easily localised within the ultrastructural images. Adding super-resolution fluorescence imaging to any correlative workflow significantly reduces the resolution gap between the fluorescence and ultrastructural studies and improves the utility of these correlative images. The ability of both 3D-SIM and SXT to image through whole tissue culture cells or tissue slices mean these two techniques are an ideal combination for rapid collection of multi-micron 3D correlative image stacks. While in this manuscript we describe a setup that combines 3D-SIM and SXT, cryoSIM is entirely compatible with a variety cryogenic ultrastructural imaging techniques, including electron microscopy methods.

## Supporting information

Custom parts diagrams

## Acknowledgements

We are grateful to Micron Oxford and its numerous partners and staff for discussions and providing the environment required for the success of complex interdisciplinary technology development project and the staff at the Diamond Light Source beamline B24 for user-support and many helpful discussions during the development of this instrument and writing this paper.

## Author contributions

Original concept - JWS, ID, IMD

Project management - MH, IMD, MJB, ID, DIS

Hardware design and construction - MAP, IMD, MJB

Software development - MAP, DMSP, IMD

Instrument commissioning - MAP, MH, IMD

Sample preparation and imaging - MH, RMP, AC, AP, MGM, IMD, IK

Writing of manuscript - IMD, ID, MJB

## Disclosures

The authors declare no conflicts of interest.

See Supplement 1 for supporting content.

