## Supplemental Figures and Table for "CryoSIM: super resolution 3D structured illumination cryogenic fluorescence microscopy for correlated ultra-structural imaging"

*Compiled May 28, 2020*

**This document provides supplementary information to "CryoSIM: super resolution 3D structured illumination cryogenic fluorescence microscopy for correlated ultra-structural imaging,"**

### 1. SUPPLEMENTAL FIGURES

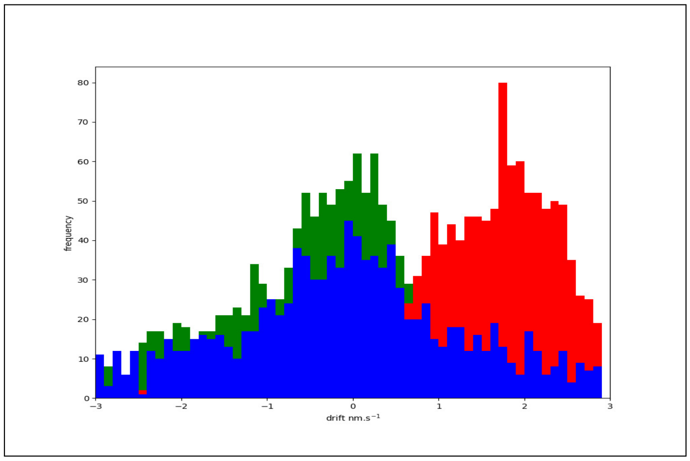


**Fig. S2.** Histogram of system drift over time: Drift rates in X (red), Y(green), and Z(blue), showing that Y and Z are roughly Gaussian distributions around 0 nm.s$^{-1}$ while the X direction has a distribution around 2 nm.s$^{-1}$ due to the rapid fill and slow emptying of the on stage dewar.

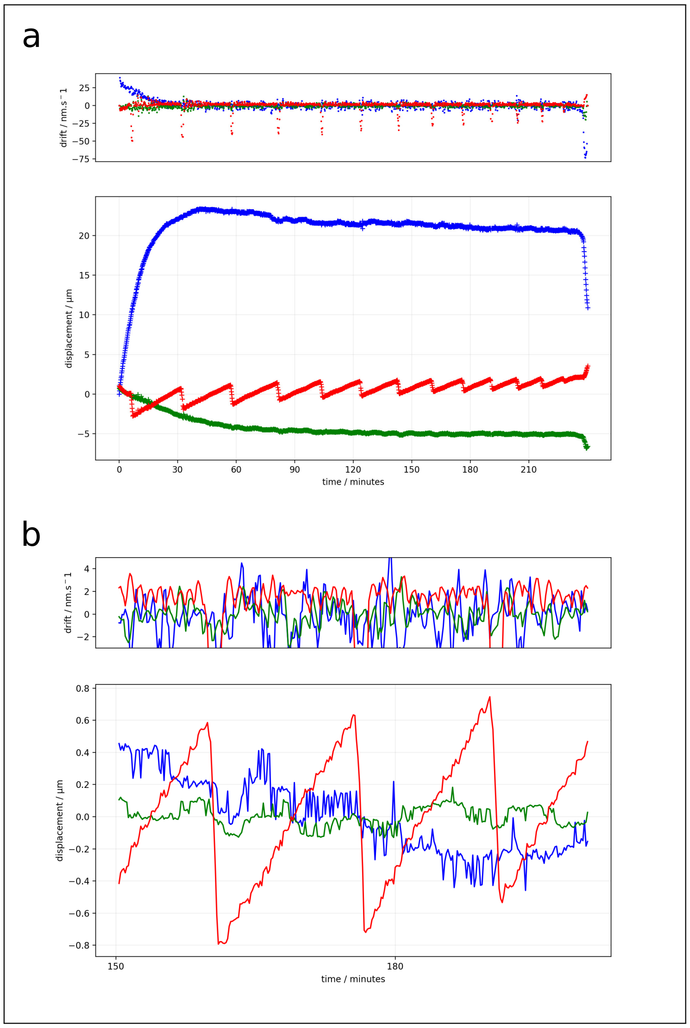


**Fig. S1.** System drift over time: Drift measured from bead centroids in 3D image stacks. (**a**) Drift rate and total drift over 4 hours in X (red), Y (green) and Z (blue). There are large changes in displacement for the first 30 minutes, especially in Z as the stage cools from room temperature down to cryogenic temperatures. Then there is a cyclic change as the on stage liquid nitrogen dewar is filled rapidly and empties over about 20 minutes. (**b**) The same data plotted from 150-200 minutes. It can be seen that the drift is in nm/s range with the x-axis having about 2 nm/s positive drift between refills. Mean displacements are fairly random about 0.2 $\mu$m in z and 0.1 $\mu$m in y over a cycle, while x undergoes a repeatable, slow drift of about 1.4 $\mu$m.

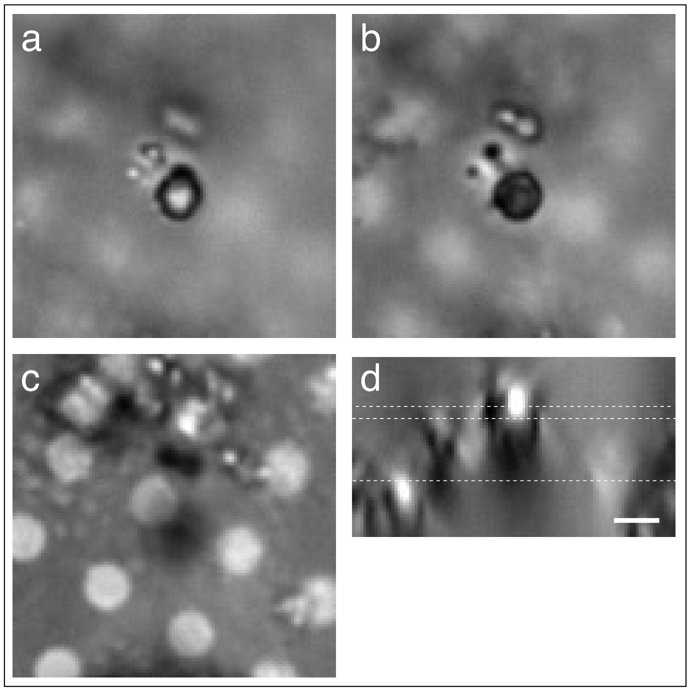


**Fig. S3.** Measurement of ice thickness above the sample: Transmission light images of plunge frozen U2OS cells. (**a**) An in focus ice crystal on top of the vitreous sample.(**b**) The same region with the focus moved down until the first sample features, 2 black vesicles just next to the ice, are in focus. (**c**) The same region again, with the carbon film supporting the sample in focus. (**d**) A YZ image through the sample at the ice crystal, with the position of the infocus plane on the ice crystal, marked at the top, 2.625 $\mu$m above the carbon, the first sample features, marked in the middle, 1.875 $\mu$m above the carbon, and the in focus carbon film at the bottom, marked with dashed white lines. The ice thickness must be less than 0.75 $\mu$m. Scale bar 2 $\mu$m.

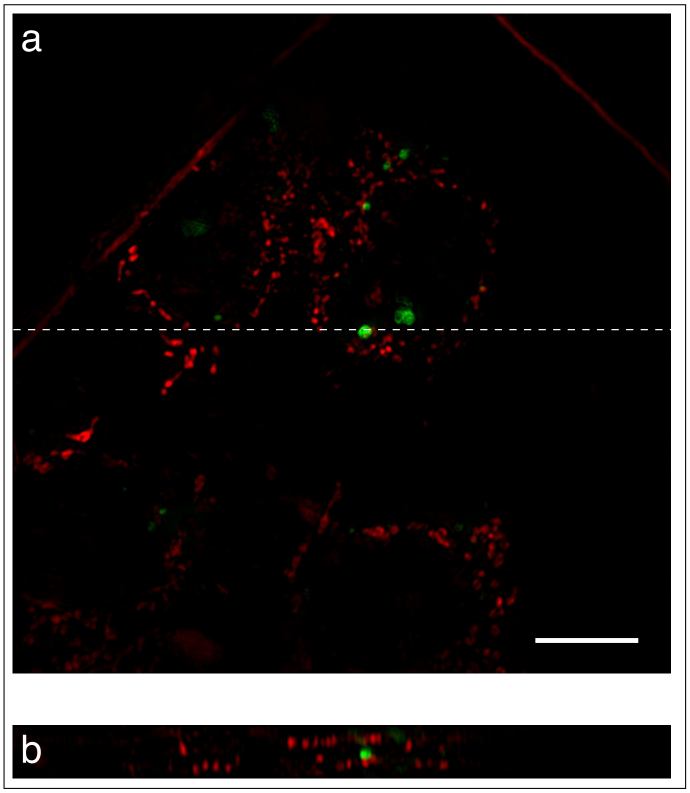

**Fig. S4.** XY and XZ slices from Fig. 6: HeLa cells expressing mov10-YFP and labelled with MitoTracker Deep Red. (**a**) SIM reconstruction fluorescence image from A maximum intensity projection over 3.125 $\mu$m with mov10-YFP in green and MitoTracker Deep Red in red. (**b**) XZ slice though the 3D data stack along the dashed line in **a**. Scale bar 10 $\mu$m.

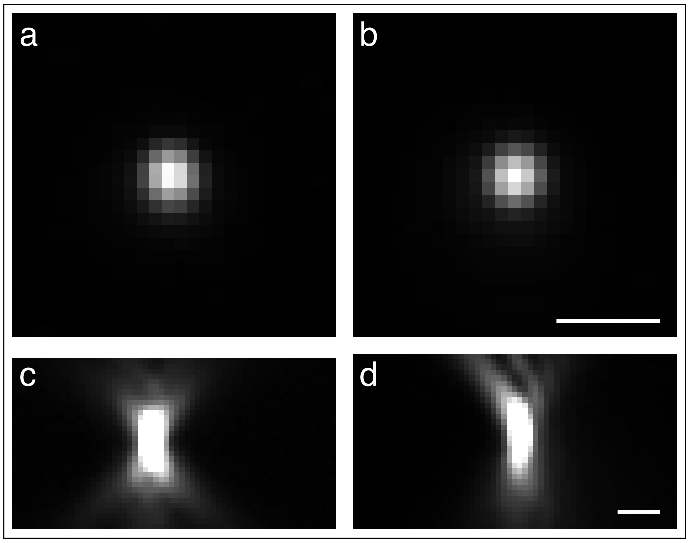

**Fig. S5.** Room temperature and Cryo PSF: Single 175 nm fluorescent beads illuminated at 488 nm and observed at with a 544/24 nm emission filter. All images are widefield. (**a,b**) in focus XY views at room and cryo temperatures. (**c,d**) XZ slices of the 3D image stack at room and cryo temperatures. These images have a lower maximum display threshold to clearly show the out of focus aberrated point spread function at cryo temperatures in **d**. Scale bars 1 $\mu$m.

### 2. TECHNICAL DRAWINGS FOR CUSTOM PARTS

The CryoSIM instrument is almost entirely constructed from commodity parts. There are a few specialist parts such as the cryo-stage and the custom made dichroic, however these parts are also commercially available. There are a small number of custom made, or adapted pieces. These are involved in the objective lifer mechanism and in cryostage stage mount which includes both precision rails for sliding the sample into and out of the optical path, and a manual coarse focus mechanism. These parts are detailed in at attached technical drawings and CAD files of them are also available on request. The following table has a list of the files and what they relate to.

**Table S1.** Technical Drawings for custom parts

| File | Description |
|---|---|
| ObjectiveMotion.pdf | Objective lifter mechanism full assembly, and diagrams of custom parts and part modifications required. |
| LinkamCarriage.pdf | Base plate for stage to run on rails. |
| ZActuator.pdf | Manual rotation wheel turns lead screw. |
| Plate.PiezoToLeadscrew.pdf | Plate under the Piezo stage that runs on the lead screw. |
| Plate.PiezoToCryostage.pdf | Adaptor plate from piezo to mount Linkam stage on. |
| BushingSpacer.pdf | Spacer to allow dual bushings and prevent the stage twisting and binding on bushings. |
| DewarCarriage.pdf | Base plate for dewar to run on rails. |
| RollerPlungerMount.pdf | Mount for a roller plunger to allow reproducible positioning of stages on rails. |
