## Supplementary material for "CryoSIM: super resolution 3D structured illumination cryogenic fluorescence microscopy for correlated ultra-structural imaging": Custom parts diagrams: BushingSpacer.pdf

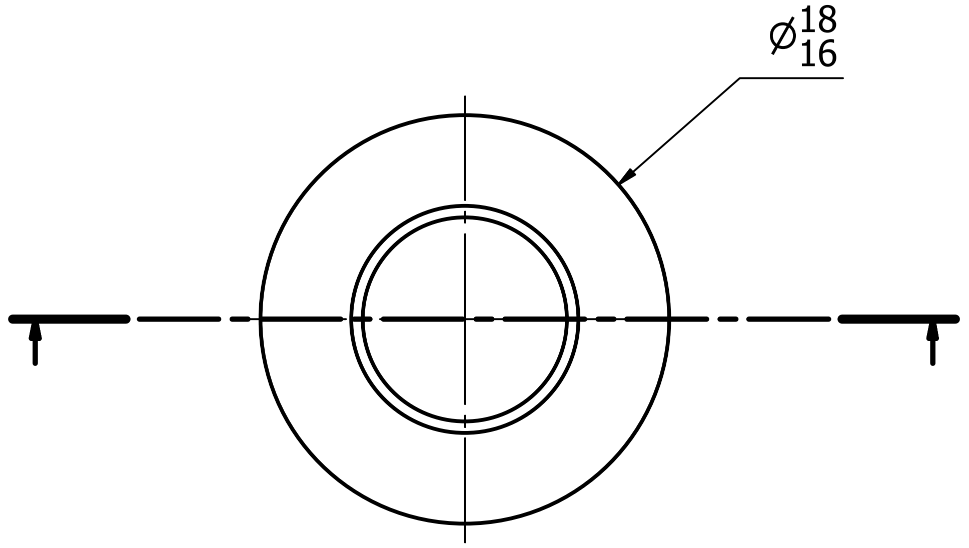


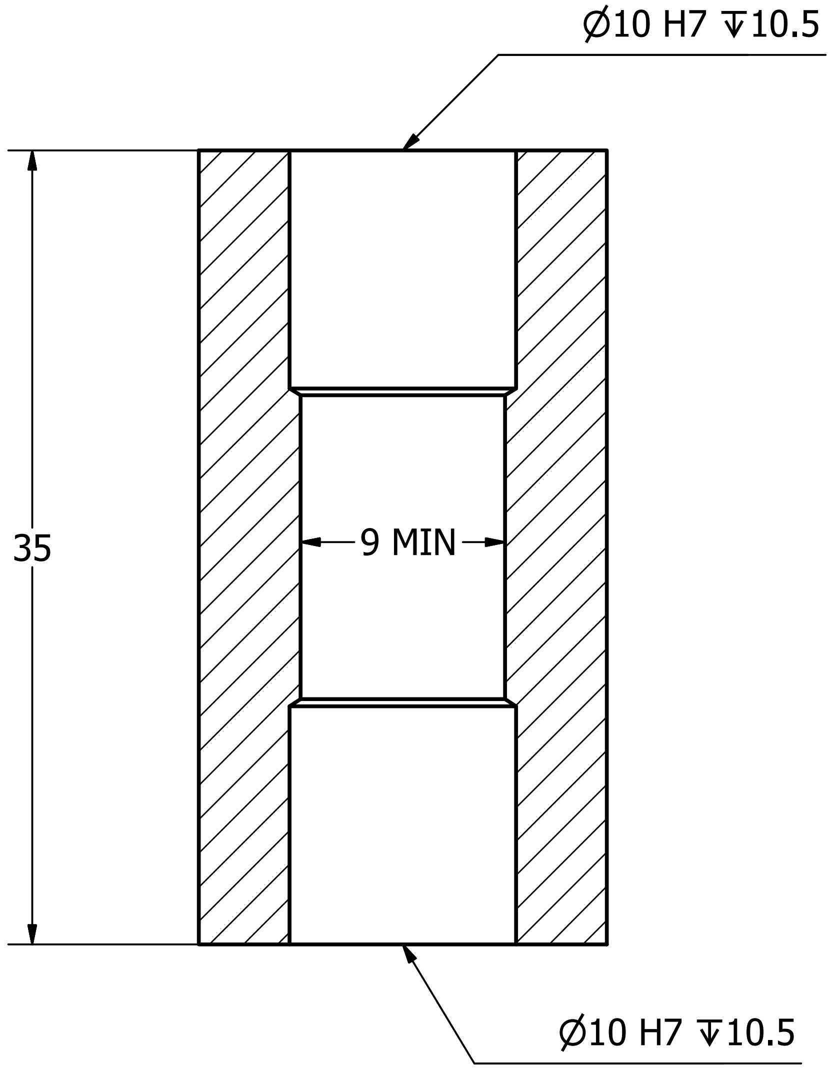


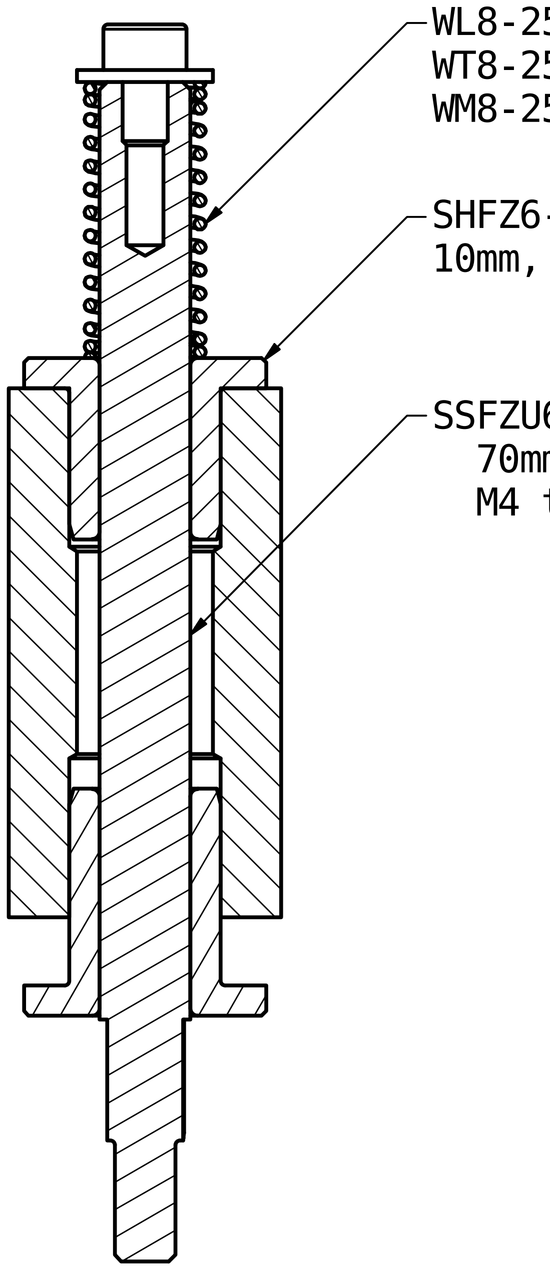


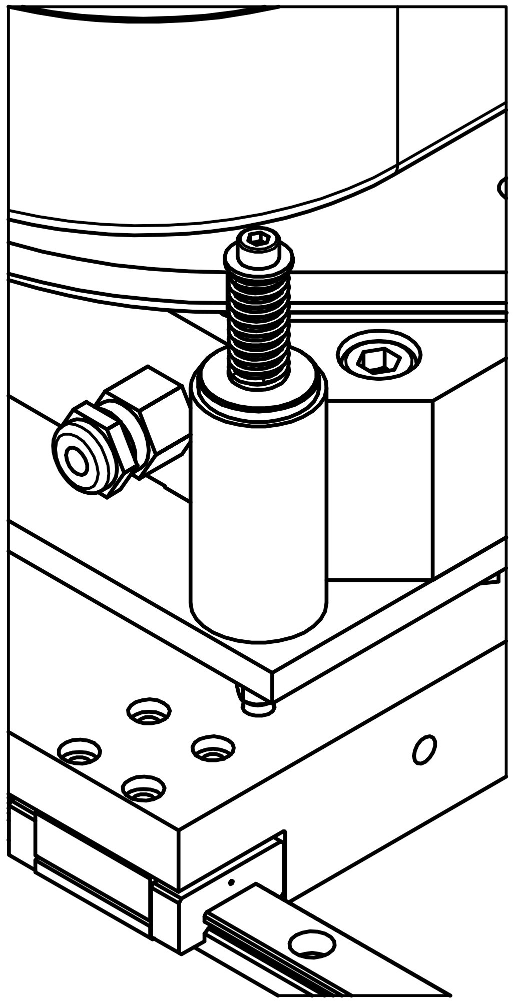

4 OFF, ALUMINIUM

| Designed by | Checked by | Approved by | Date | Date |
|---|---|---|---|---|
| map | | | | 02/11/2018 |

| | | | |
|---|---|---|---|
| 3rd ANGLE PROJECTION | Micron Oxford | | |
| | | BushingSpacer | Edition | Sheet 1 / 1 |
