## Supplementary material for "CryoSIM: super resolution 3D structured illumination cryogenic fluorescence microscopy for correlated ultra-structural imaging": Custom parts diagrams: DewarCarriage.pdf

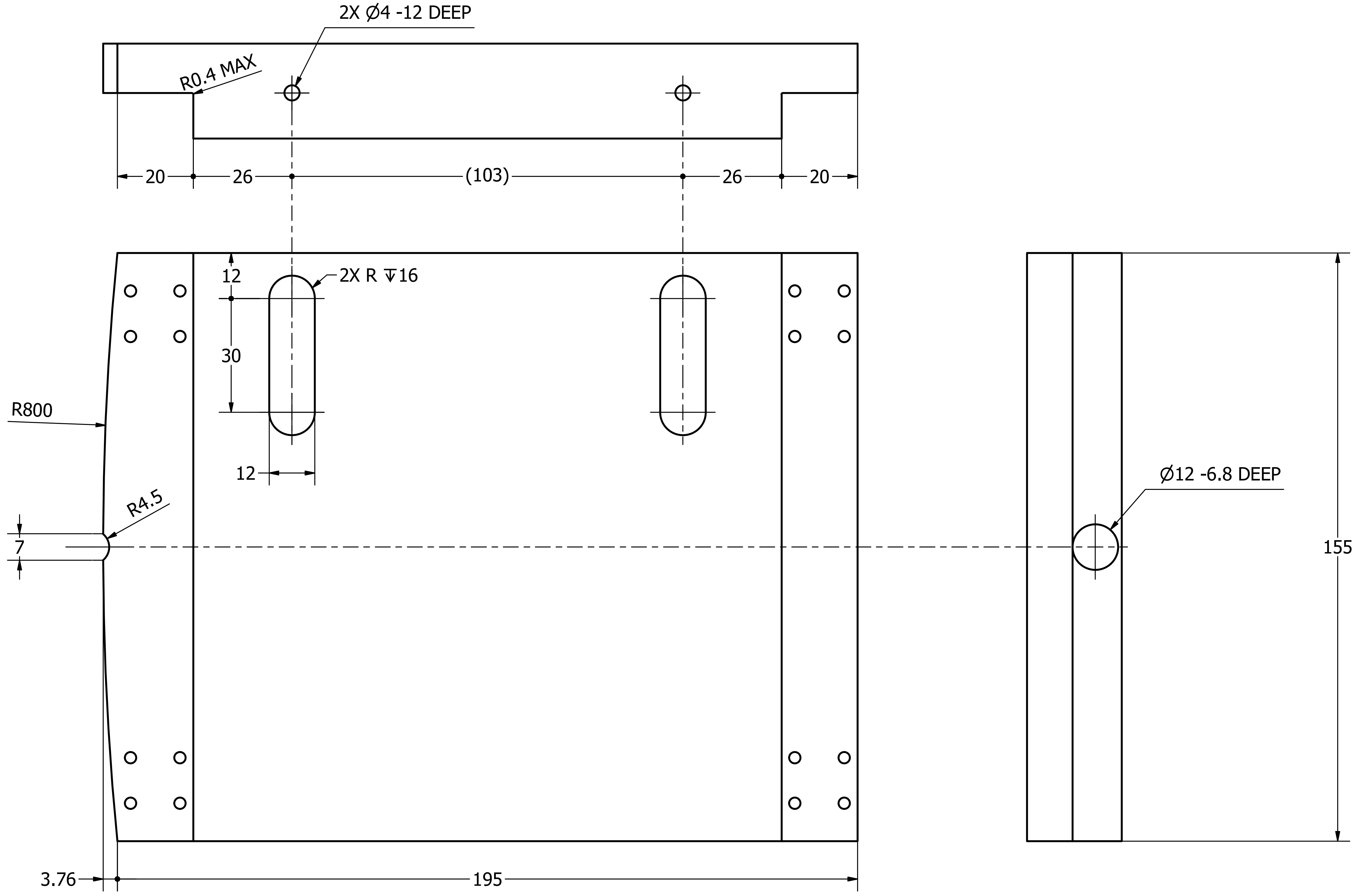


| Designed by | Checked by | Approved by | Date | Date |
|---|---|---|---|---|
| map | | | | 06/06/2019 |

3rd ANGLE PROJECTION

Micron Oxford

| DewarCarriage | Edition | Sheet |
|---|---|---|
| | | 1 / 2 |

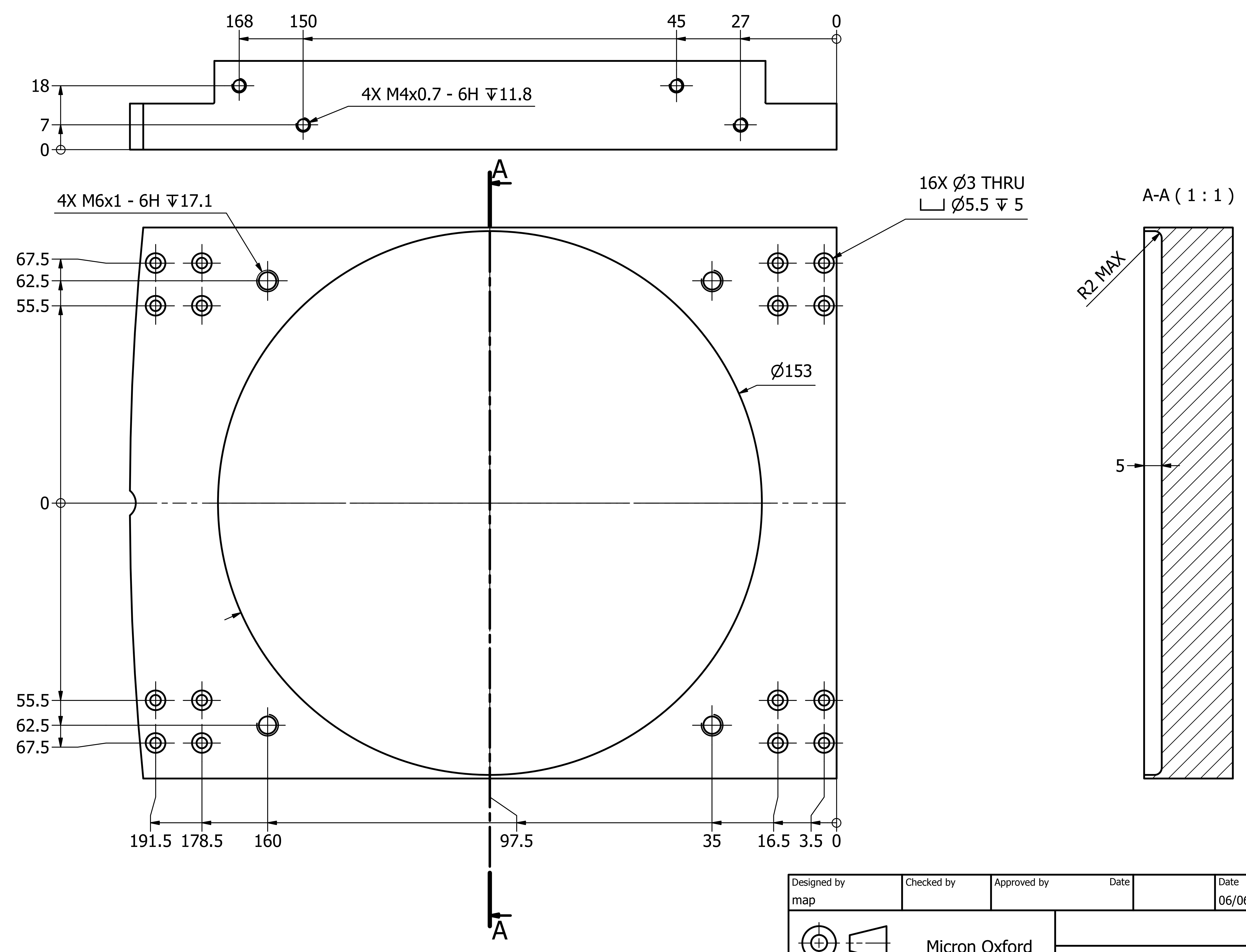


| Designed by | Checked by | Approved by | Date | Date |
|---|---|---|---|---|
| map | | | | 06/06/2019 |

3rd ANGLE PROJECTION

Micron Oxford

| | Edition | Sheet |
|---|---|---|
| DewarCarriage | | 2 / 2 |
