## Supplementary material for "CryoSIM: super resolution 3D structured illumination cryogenic fluorescence microscopy for correlated ultra-structural imaging": Custom parts diagrams: LinkamCarriage.pdf

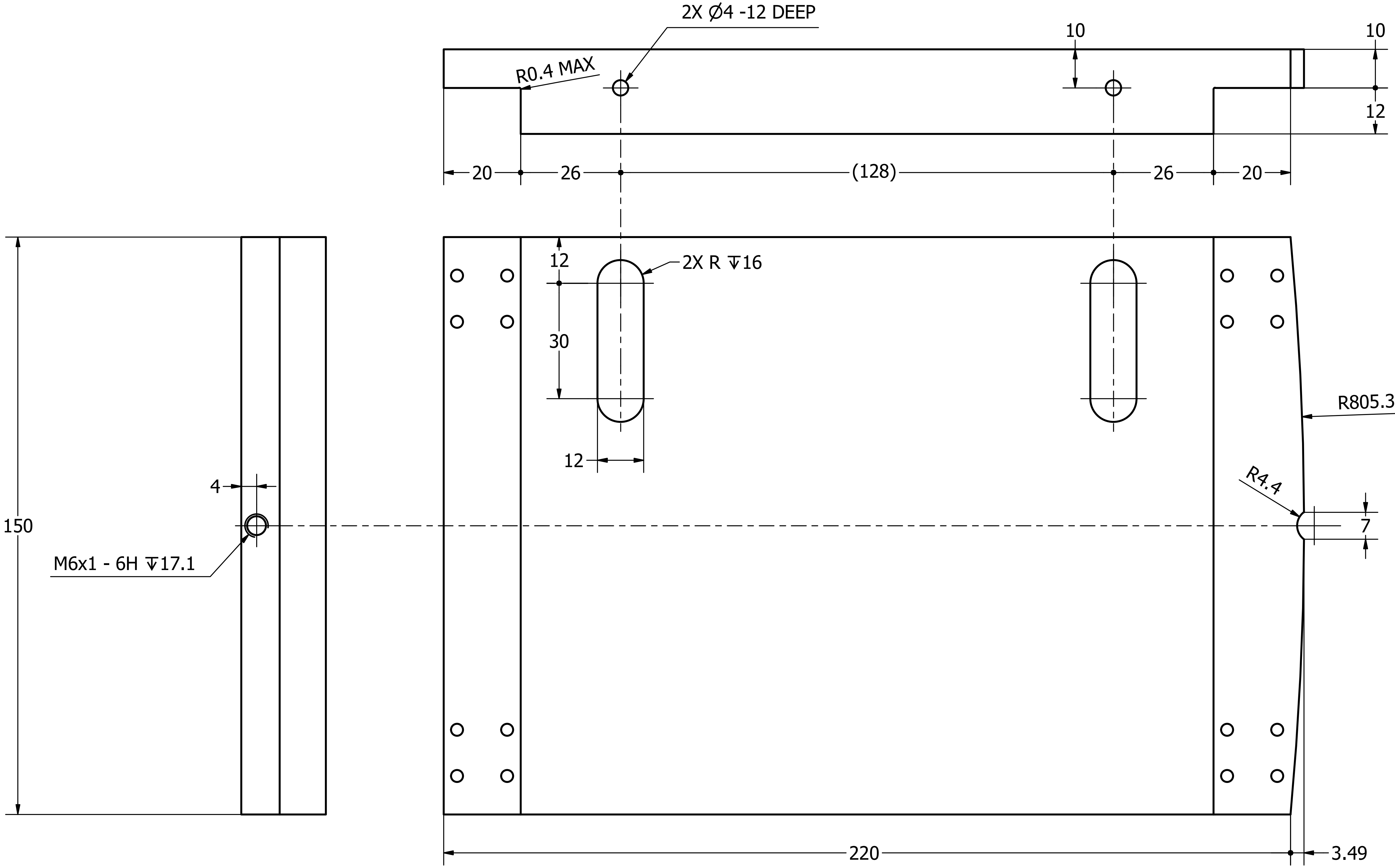

1 OFF, ALUMINIUM

| Designed by | Checked by | Approved by | Date | Date |
|---|---|---|---|---|
| map | | | | 14/08/2018 |

3rd ANGLE PROJECTION

Micron Oxford

| LinkamCarriage | Edition | Sheet |
|---|---|---|
| | | 1 / 2 |

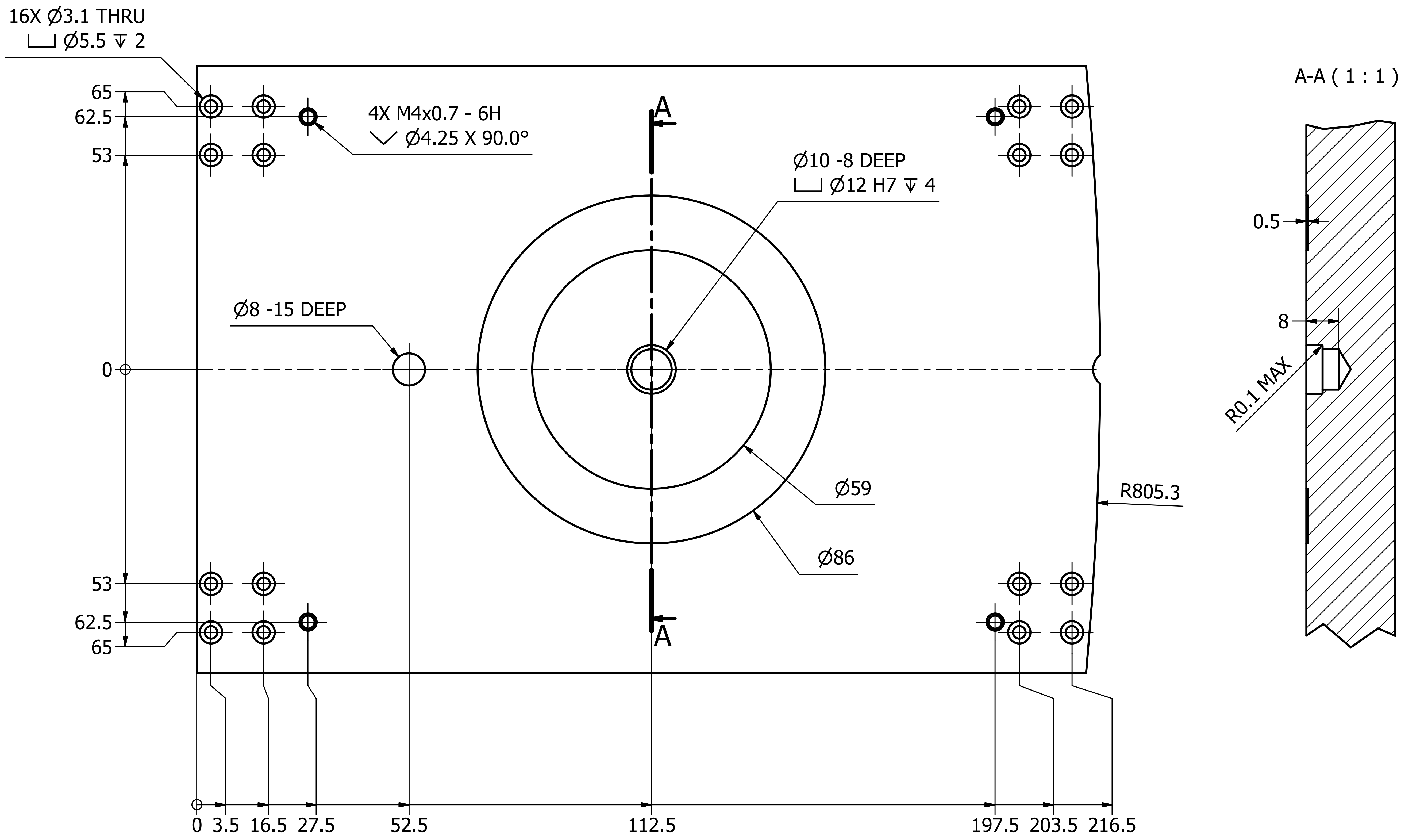

| Designed by | Checked by | Approved by | Date | Date |
|---|---|---|---|---|
| map | | | | 14/08/2018 |

| 3rd ANGLE PROJECTION | Micron Oxford | LinkamCarriage | Edition | Sheet |
|---|---|---|---|---|
| | | | | 2 / 2 |
