## Supplementary material for "CryoSIM: super resolution 3D structured illumination cryogenic fluorescence microscopy for correlated ultra-structural imaging": Custom parts diagrams: ObjectiveMotion.pdf

### mirror mount ( 1 : 1 )

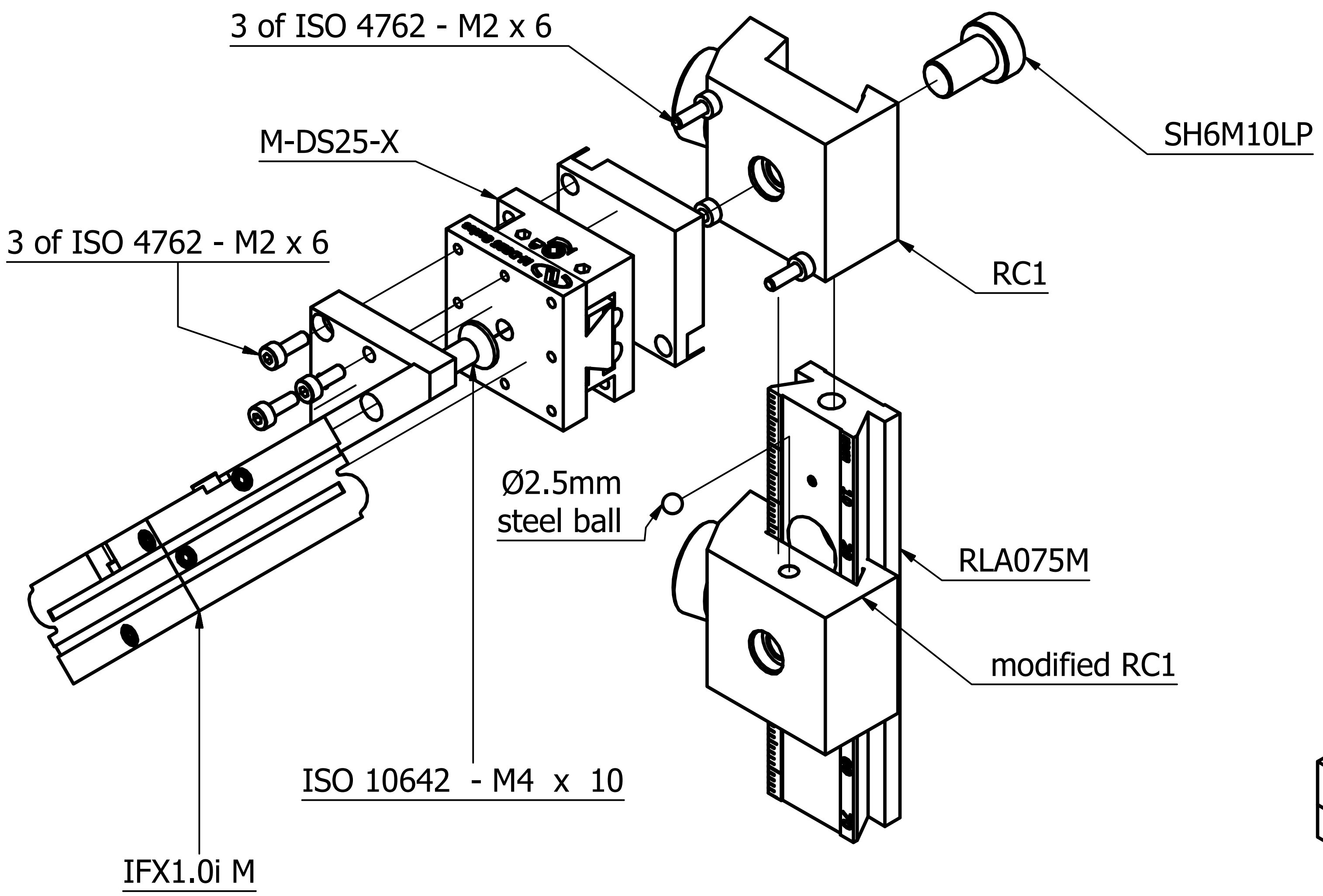

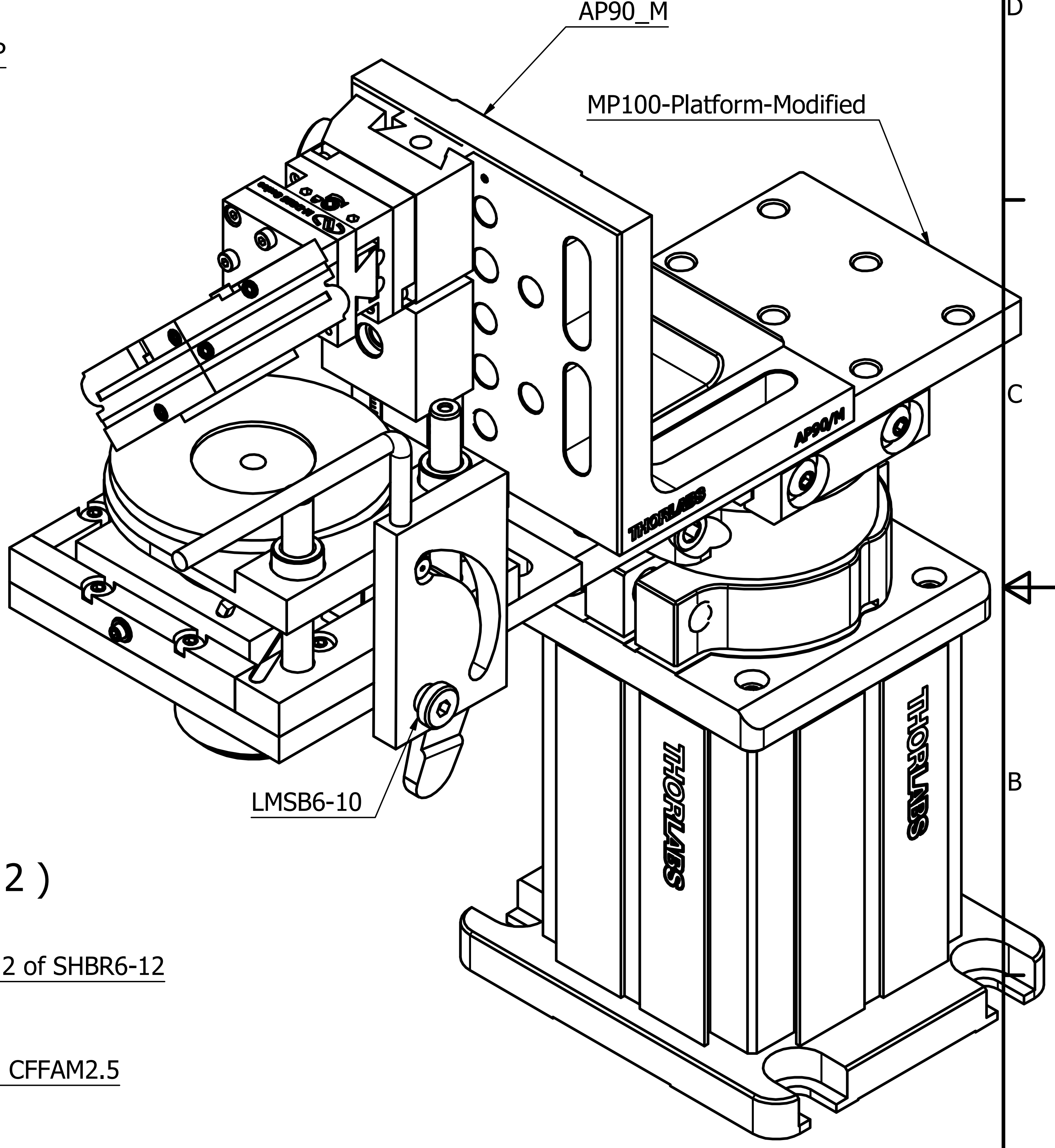

### objective mount ( 1 : 2 )

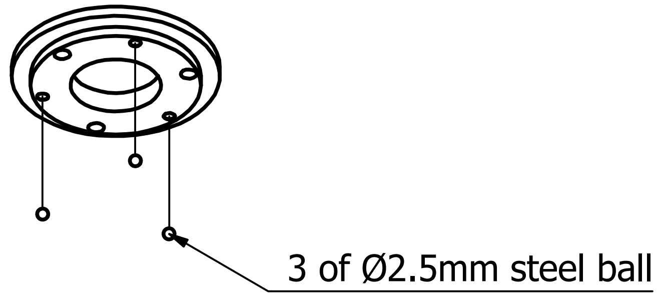

### lifter plate ( 1 : 2 )

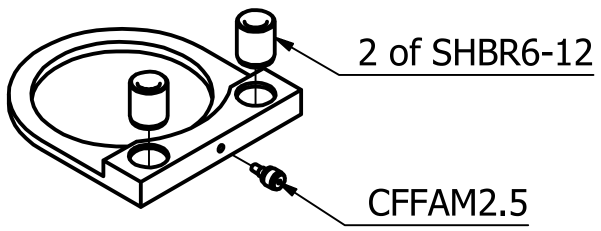

| DESIGNED BY | CHECKED BY | APPROVED BY | DATE | DATE | |
|---|---|---|---|---|---|
| map | | | | 13/05/2019 | |
| | | | B24 Objective mount | ISSUE 2 | SHEET 1 / 12 |

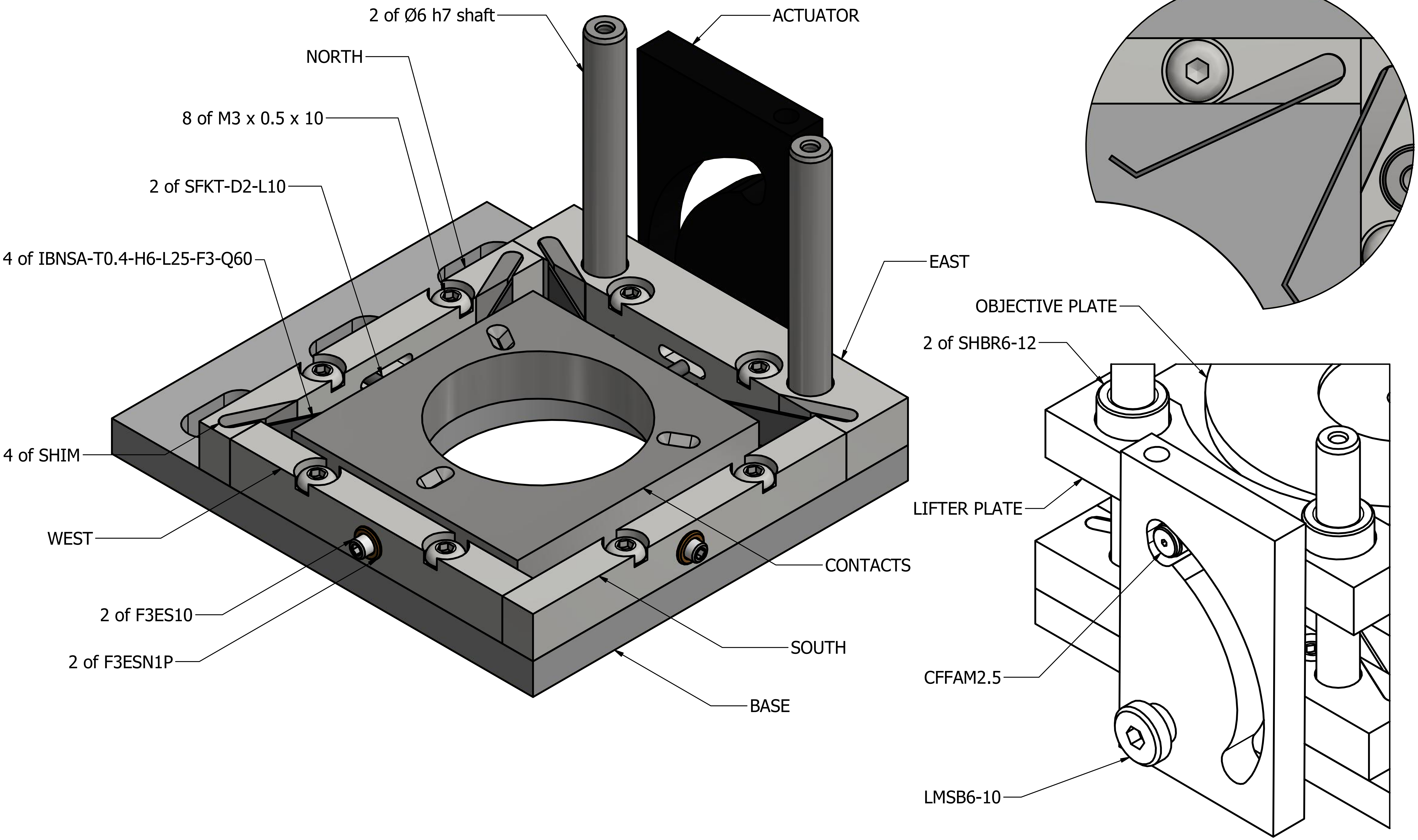

| Designed by | Checked by | Approved by | Date | | Date | |
|---|---|---|---|---|---|---|
| map | | | | | 13/05/2019 | |
| | | B24 Objective mount | | | Edition 2 | Sheet 2 / 12 |

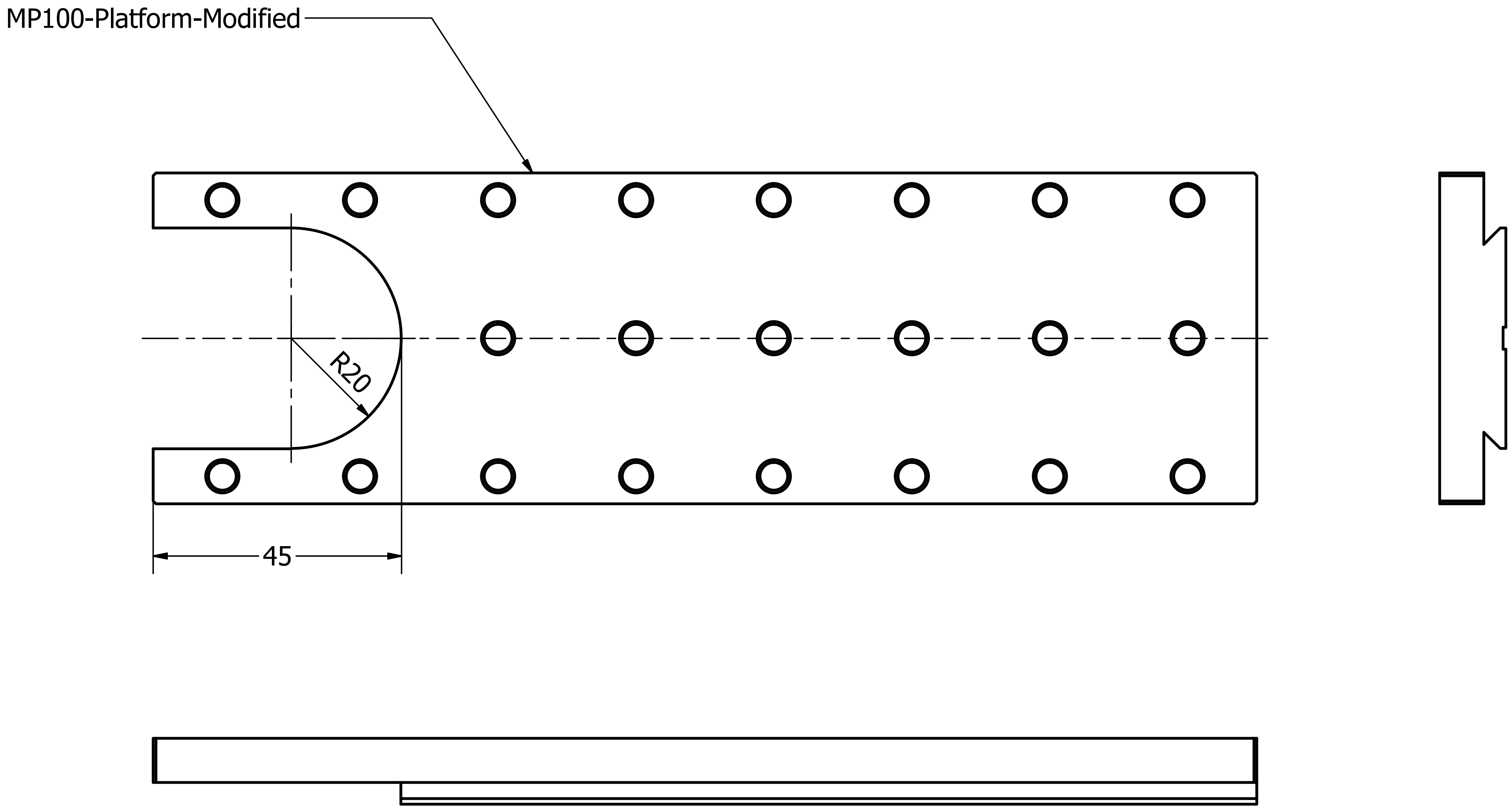

1 feature to be cut/milled.
Part supplied in aluminium.

| Designed by | Checked by | Approved by | Date | Date |
|---|---|---|---|---|
| map | | | | 13/05/2019 |

| Title | Edition | Sheet |
|---|---|---|
| B24 Objective mount | 2 | 3 / 12 |

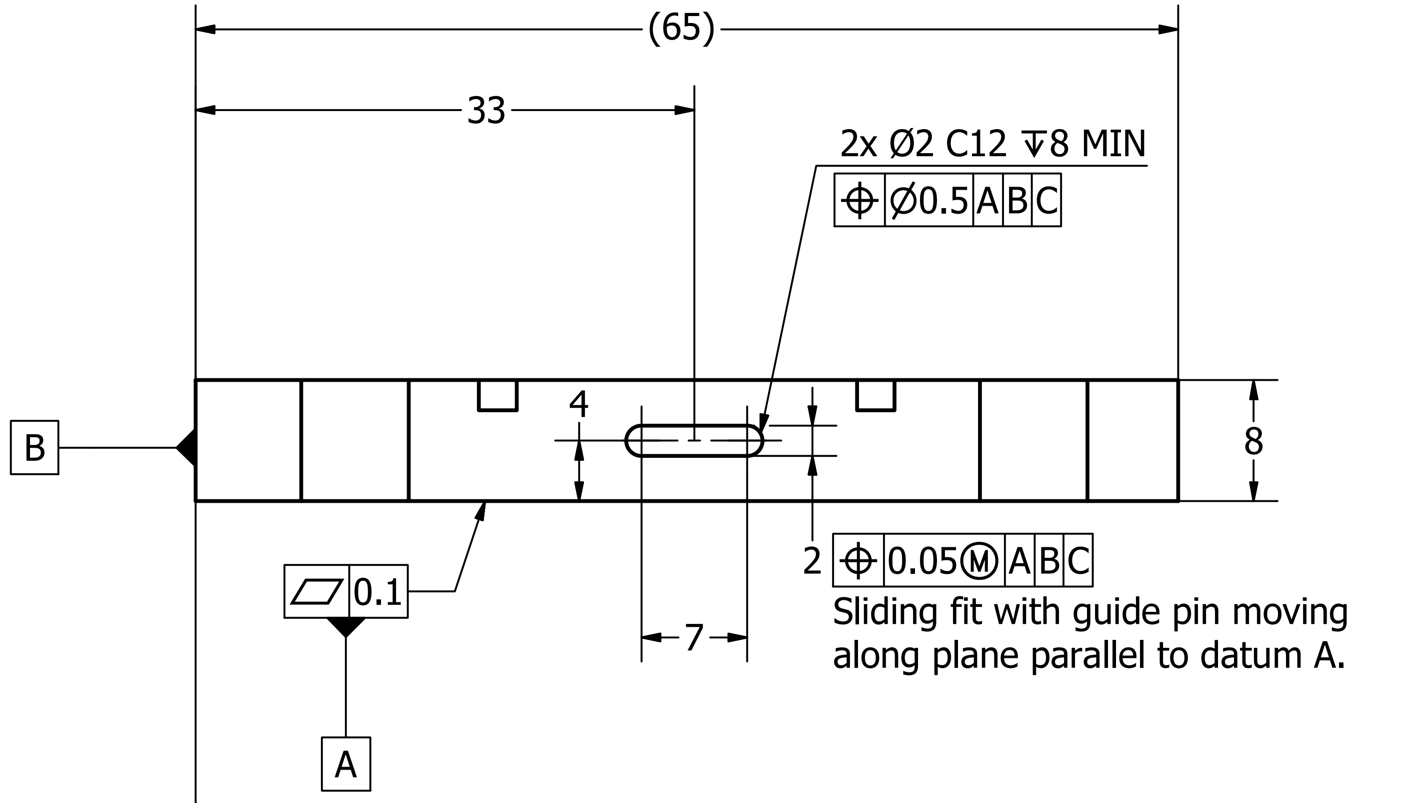

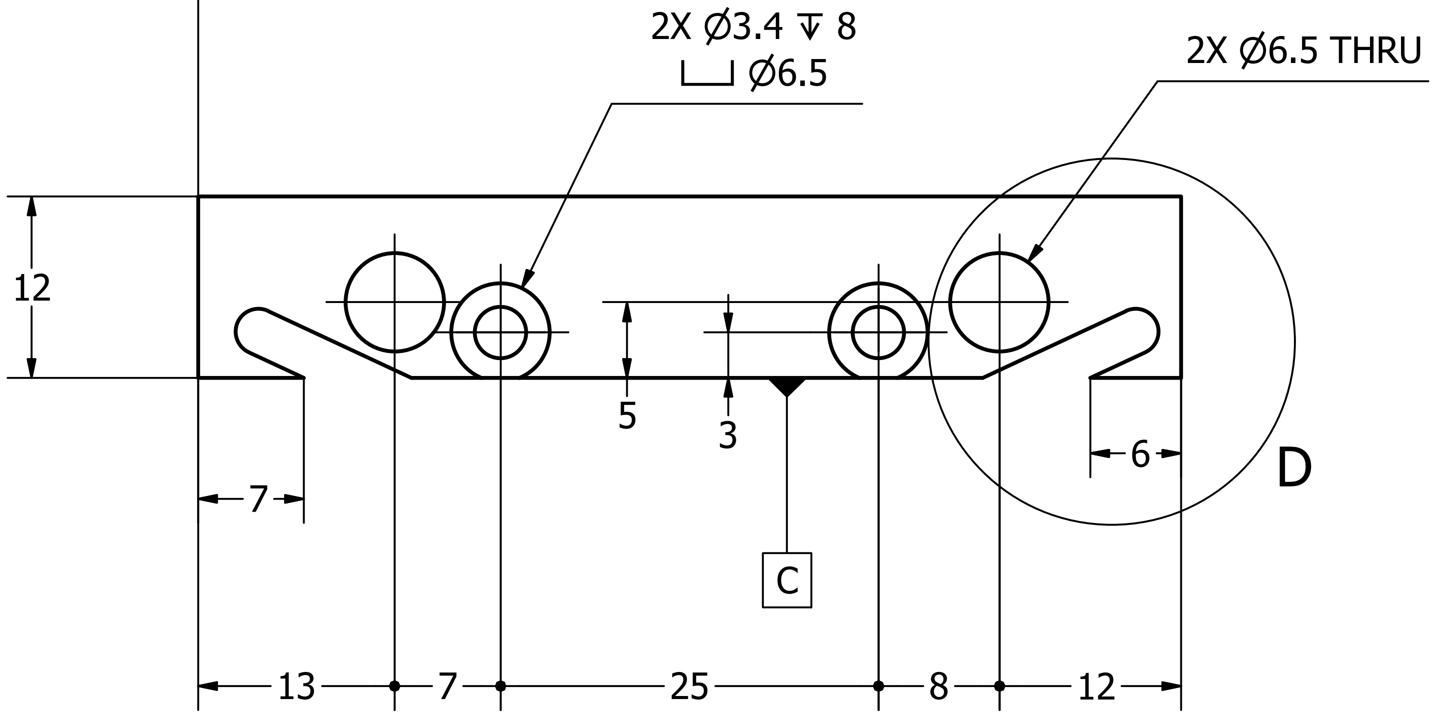

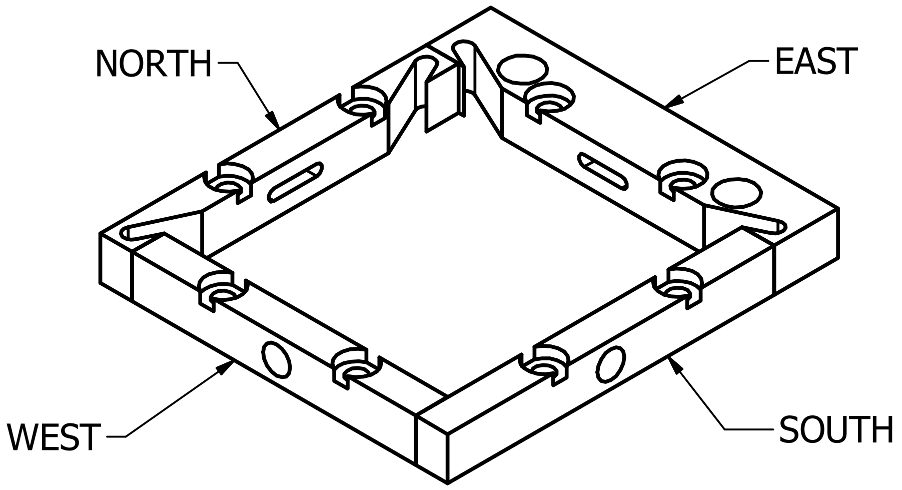

D ( 4 : 1 )

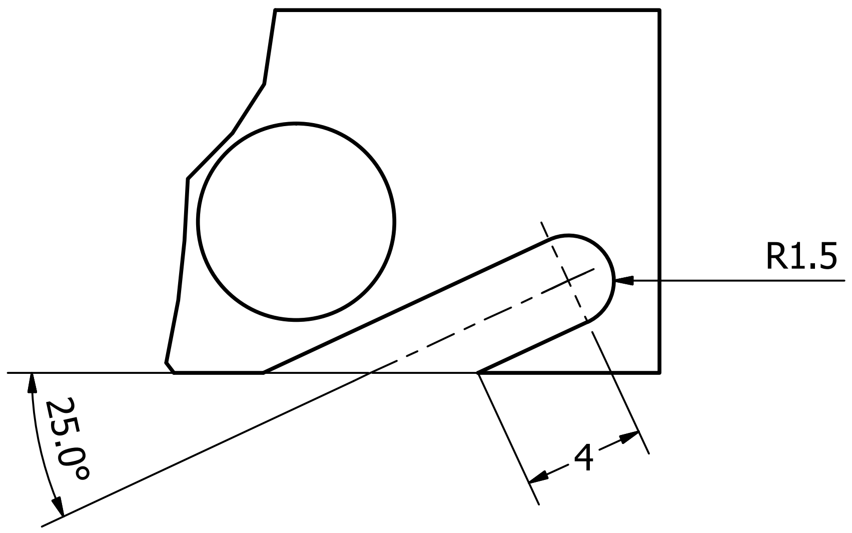

### EAST ( 2 : 1 )

1 off, steel

S275

| Designed by | Checked by | Approved by | Date | Date |
|---|---|---|---|---|
| map | | | | 13/05/2019 |

| | | |
|---|---|---|
| B24 Objective mount | Edition 2 | Sheet 4 / 12 |

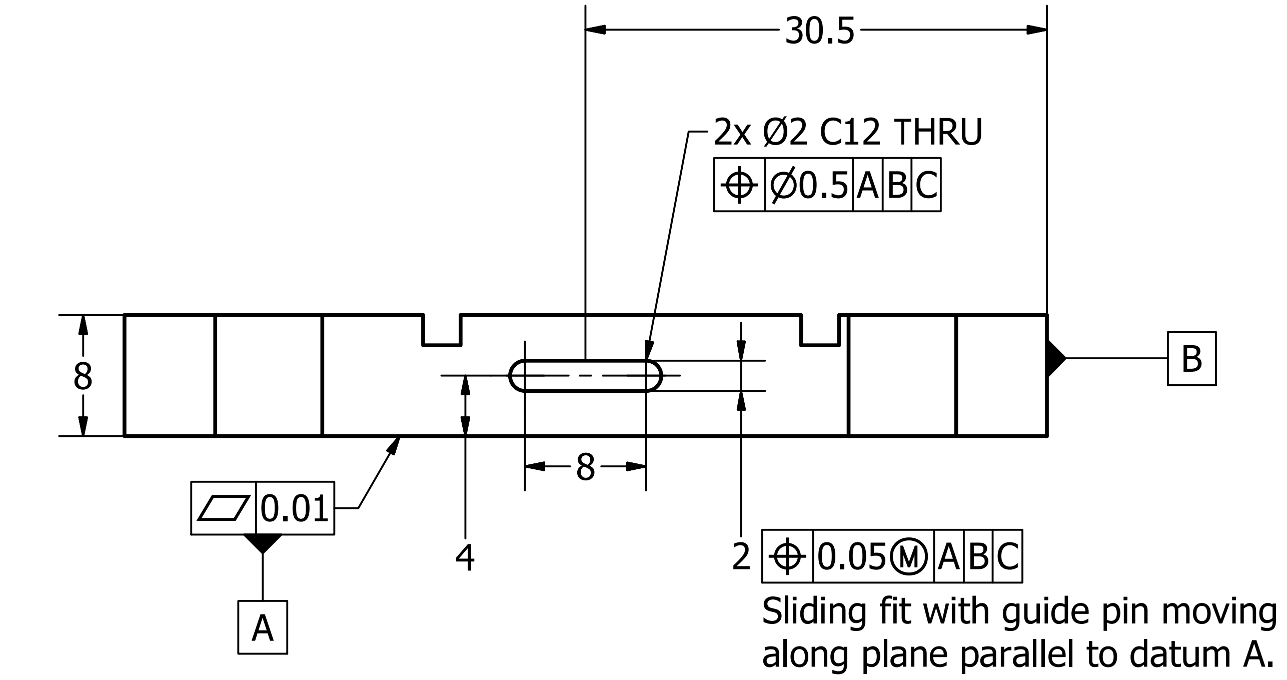

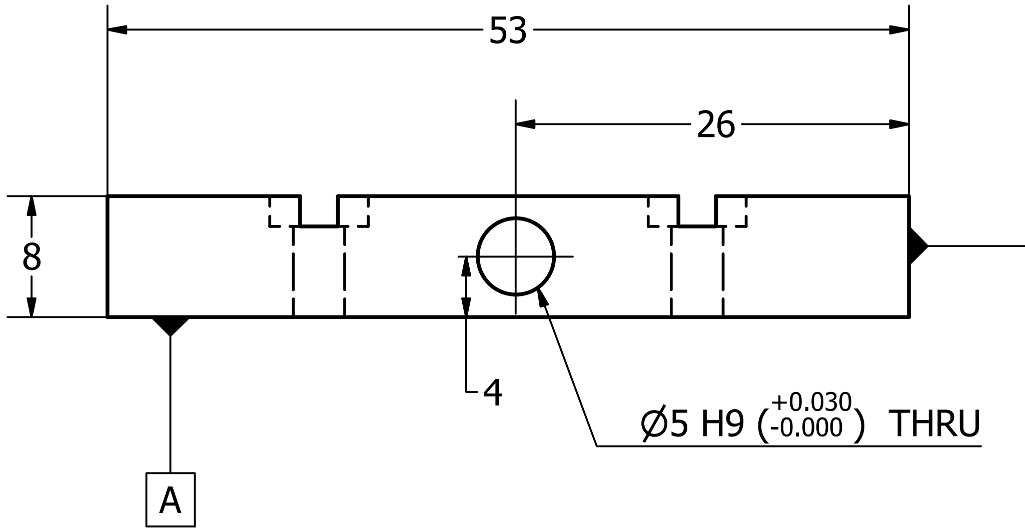

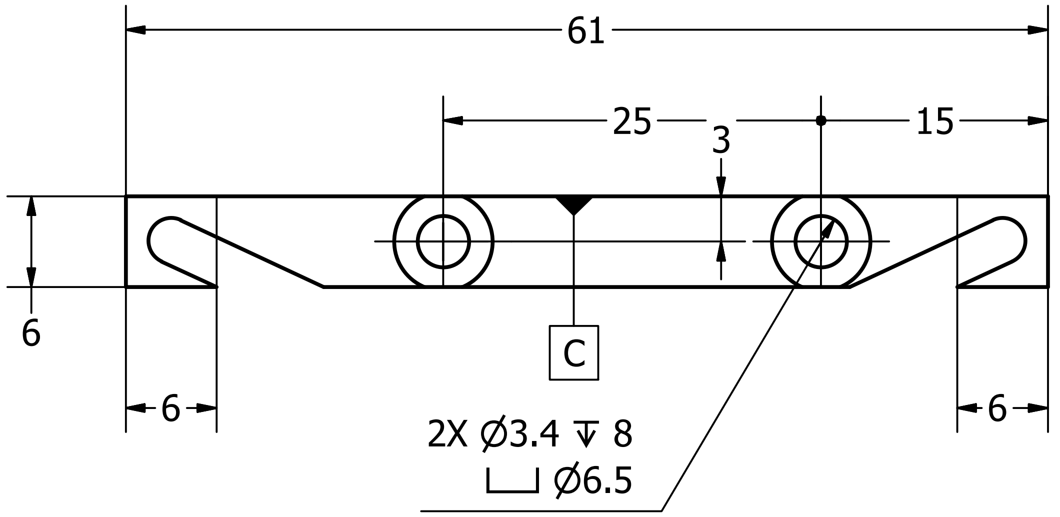

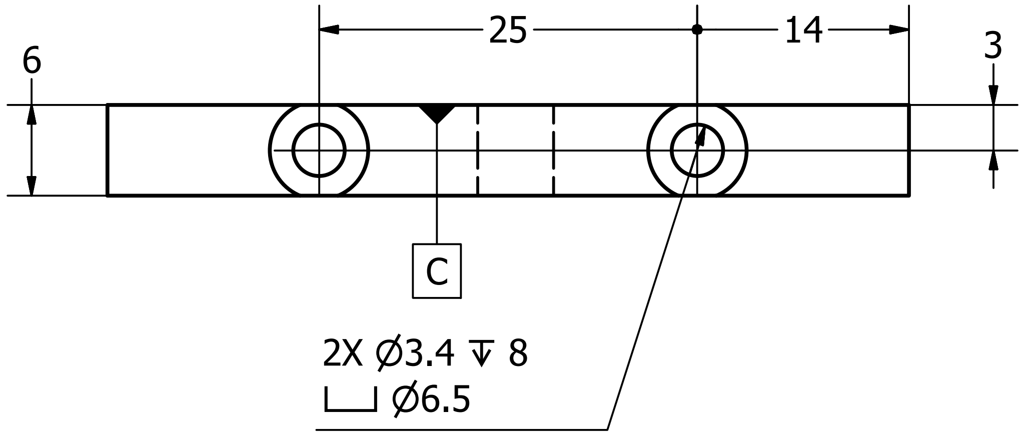

NORTH ( 2 : 1 )
1 off, steel
S275

WEST ( 2 : 1 )
1 off, steel
S275

| Designed by | Checked by | Approved by | Date | Date |
|---|---|---|---|---|
| map | | | | 13/05/2019 |

| Title | Edition | Sheet |
|---|---|---|
| B24 Objective mount | 2 | 5 / 12 |

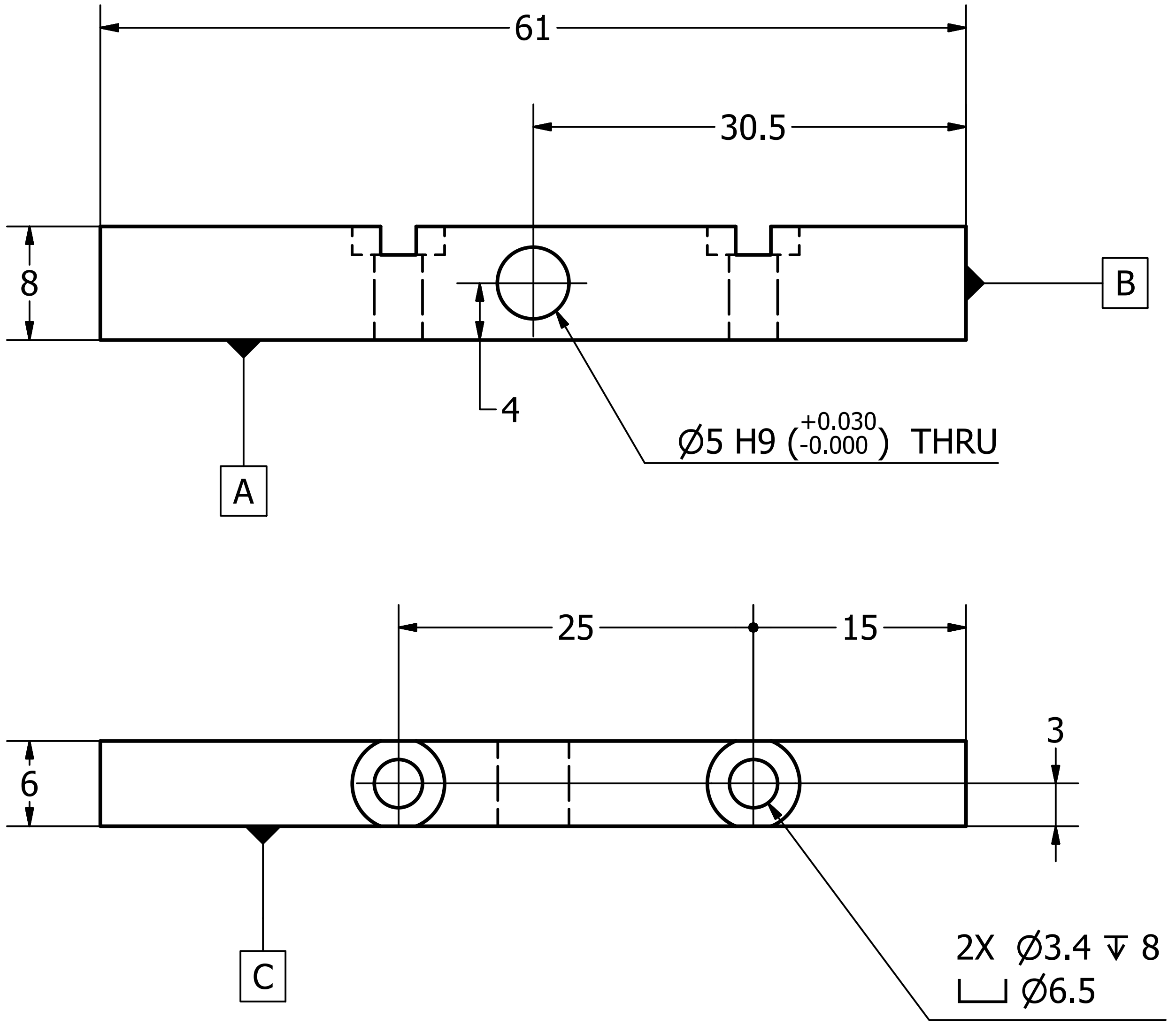

SOUTH ( 2 : 1 )
1 off, steel
S275

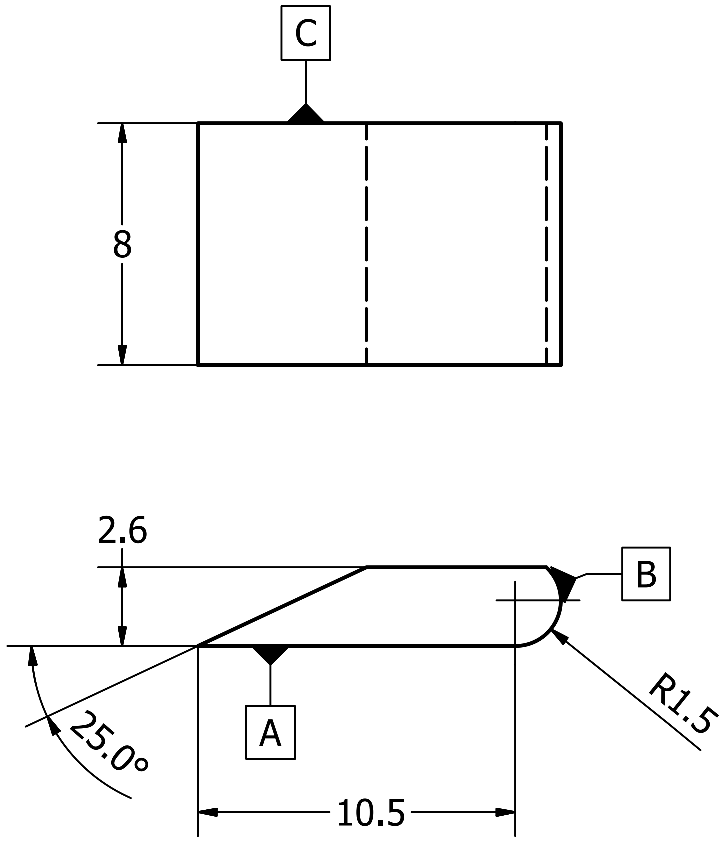

SHIM ( 4 : 1 )
4 off, steel

| Designed by | Checked by | Approved by | Date | Date | |
|---|---|---|---|---|---|
| map | | | | 13/05/2019 | |
| B24 Objective mount | | | | Edition 2 | Sheet 6 / 12 |

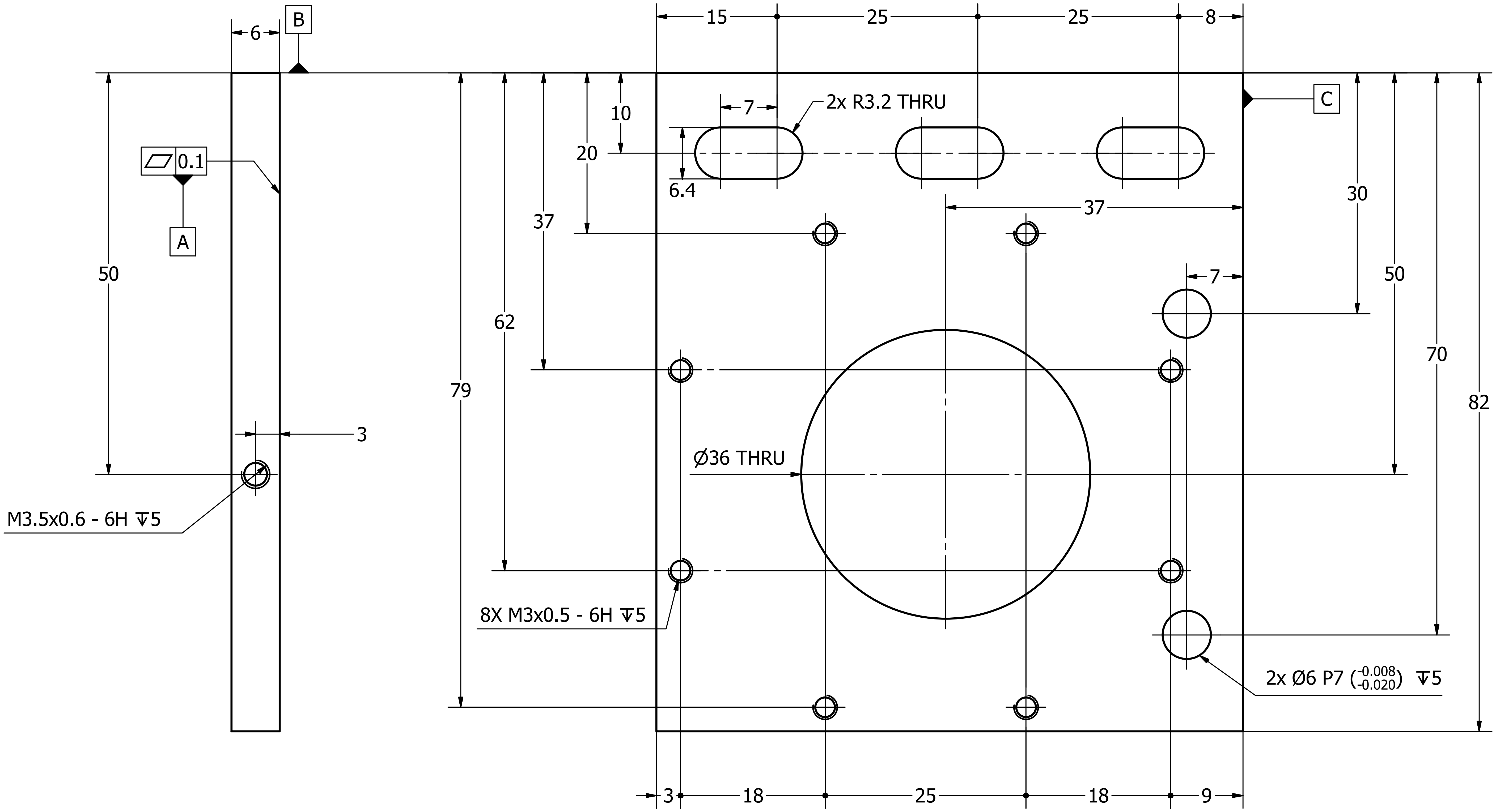

### BASE ( 2 : 1 )

1 off, steel

EN1A S275 or 6mm gauge plate

| Designed by | Checked by | Approved by | Date | Date |
|---|---|---|---|---|
| map | | | | 13/05/2019 |

| | Edition | Sheet |
|---|---|---|
| B24 Objective mount | 2 | 7 / 12 |

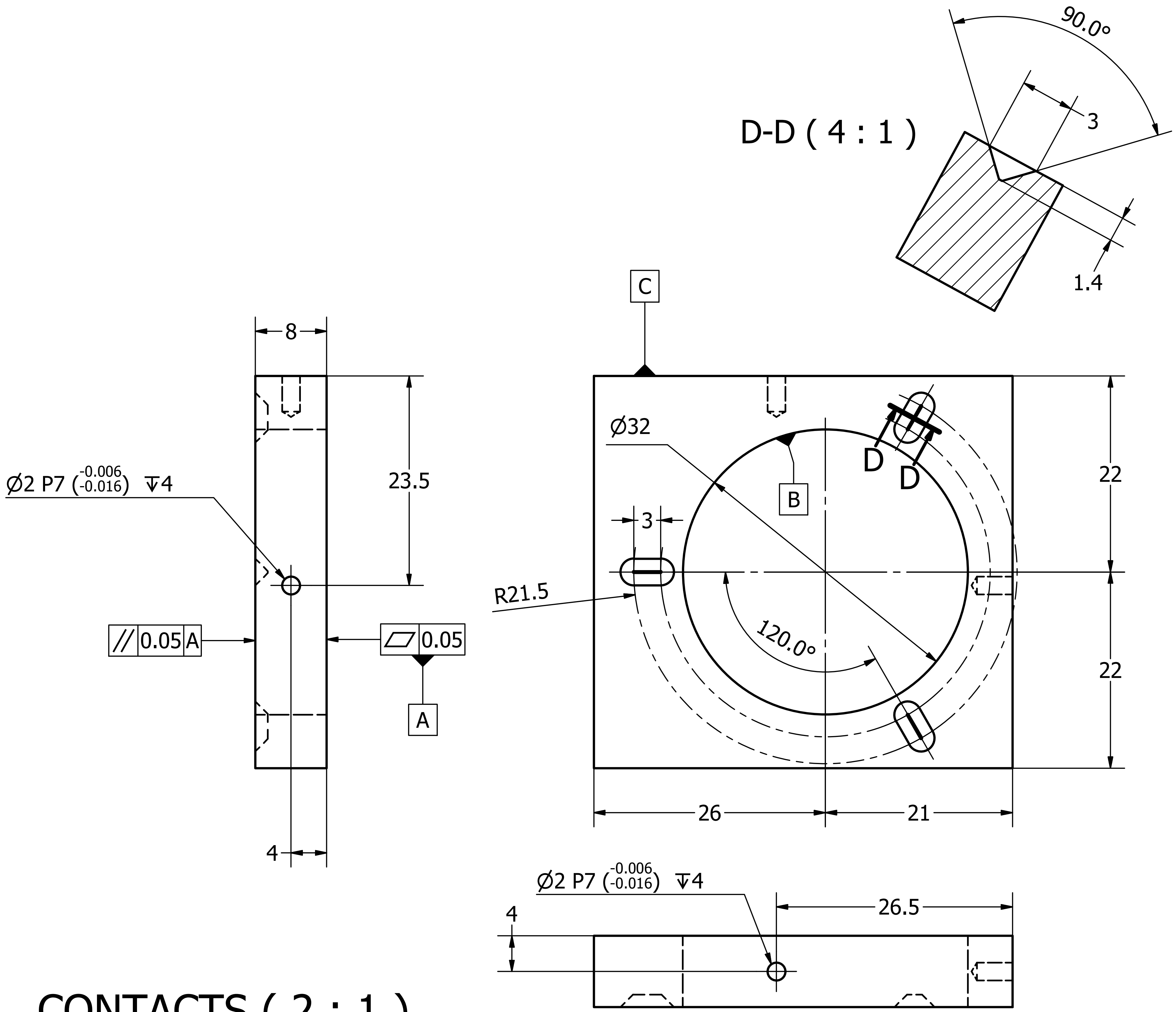

### CONTACTS ( 2 : 1 )
#### 1 off, steel
#### S275 or 8mm gauge plate

| Designed by | Checked by | Approved by | Date | Date |
|---|---|---|---|---|
| map | | | | 13/05/2019 |
| | B24 Objective mount | | Edition 2 | Sheet 8 / 12 |

CAM CL (centreline) co-ordinates along directions
AB and CB, with axis of feature D at (0,0).
(Supplied as CAM.CSV with 0.1 steps along AB.)

| CAM CL | | CAM CL | |
|---|---|---|---|
| AB | CB | AB | CB |
| 0.0 | 14.5 | 14.0 | 16.9 |
| 0.5 | 14.9 | 14.5 | 16.8 |
| 1.0 | 15.2 | 15.0 | 16.7 |
| 1.5 | 15.5 | 15.5 | 16.5 |
| 2.0 | 15.7 | 16.0 | 16.4 |
| 2.5 | 16.0 | 16.5 | 16.2 |
| 3.0 | 16.2 | 17.0 | 16.0 |
| 3.5 | 16.4 | 17.5 | 15.8 |
| 4.0 | 16.6 | 18.0 | 15.6 |
| 4.5 | 16.8 | 18.5 | 15.3 |
| 5.0 | 16.9 | 19.0 | 15.1 |
| 5.5 | 17.0 | 19.5 | 14.8 |
| 6.0 | 17.1 | 20.0 | 14.5 |
| 6.5 | 17.2 | 20.5 | 14.2 |
| 7.0 | 17.3 | 21.0 | 13.9 |
| 7.5 | 17.4 | 21.5 | 13.6 |
| 8.0 | 17.4 | 22.0 | 13.2 |
| 8.5 | 17.5 | 22.5 | 12.8 |
| 9.0 | 17.5 | 23.0 | 12.4 |
| 9.5 | 17.5 | 23.5 | 12.0 |
| 10.0 | 17.5 | 24.0 | 11.6 |
| 10.5 | 17.5 | 24.5 | 11.1 |
| 11.0 | 17.4 | 25.0 | 10.6 |
| 11.5 | 17.4 | 25.5 | 10.1 |
| 12.0 | 17.3 | 26.0 | 9.5 |
| 12.5 | 17.2 | 26.5 | 9.0 |
| 13.0 | 17.2 | 27.0 | 8.3 |
| 13.5 | 17.1 | 27.5 | 7.7 |

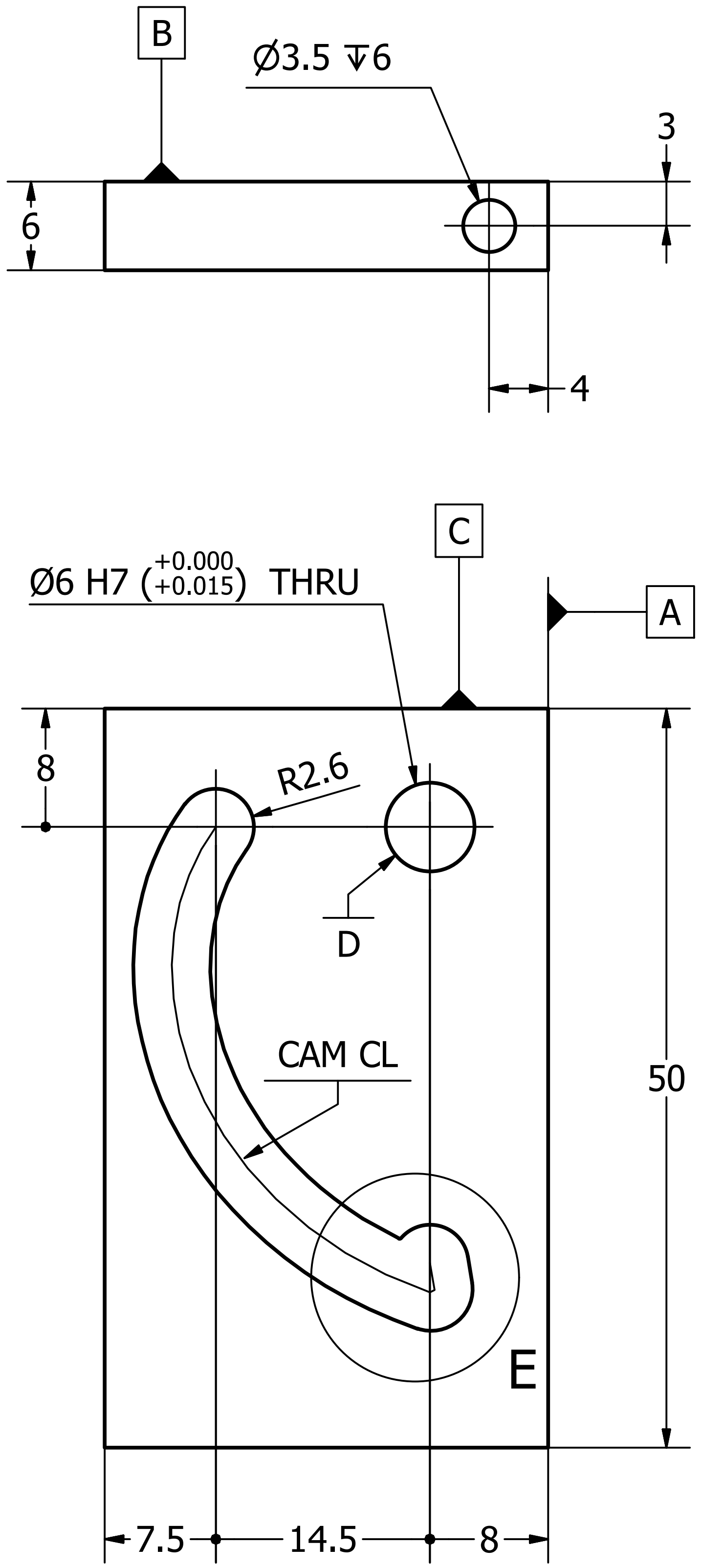

ACTUATOR ( 2 : 1 )
1 off, aluminium

E ( 10 : 1 )

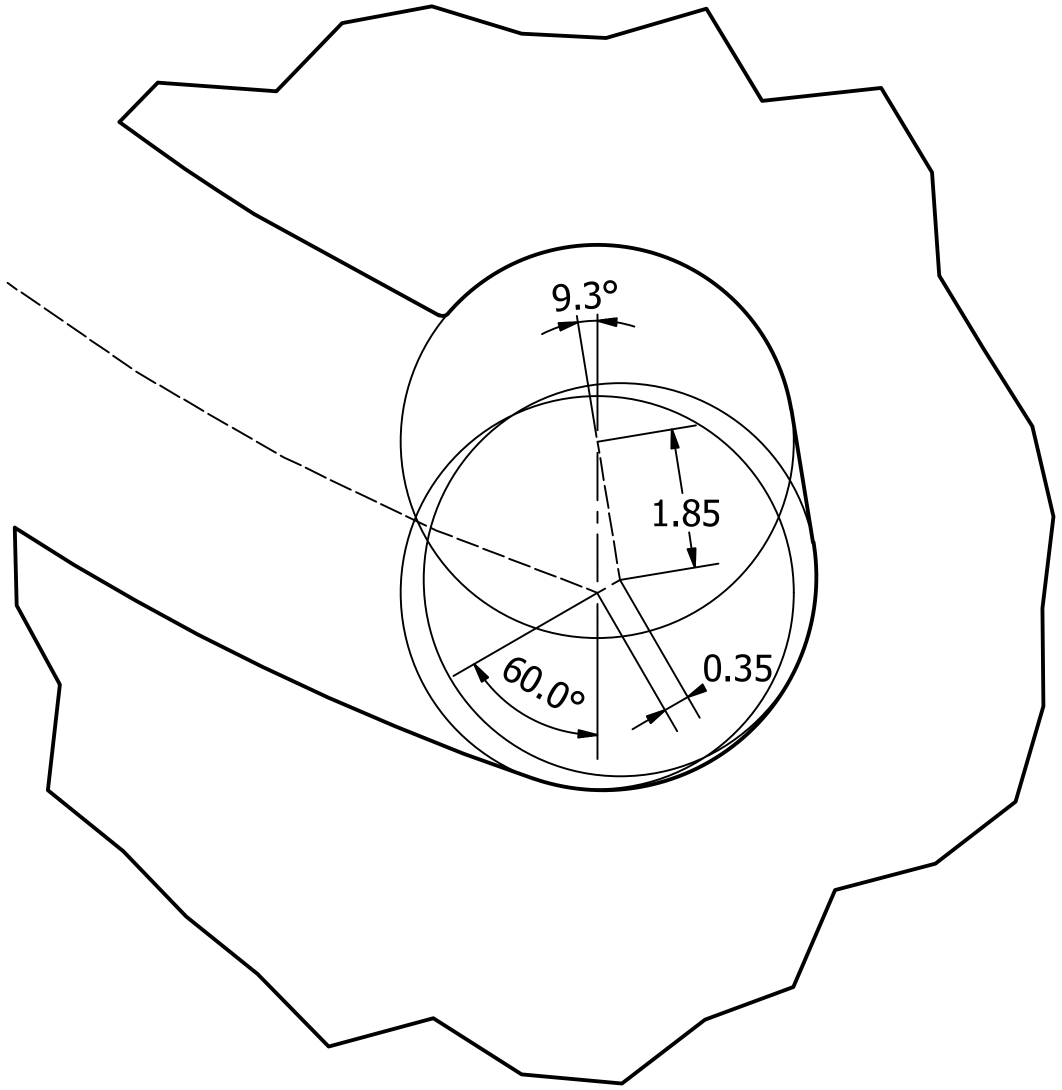

| Designed by | Checked by | Approved by | Date | Date |
|---|---|---|---|---|
| map | | | | 13/05/2019 |

| | | |
|---|---|---|
| B24 Objective mount | Edition 2 | Sheet 9 / 12 |

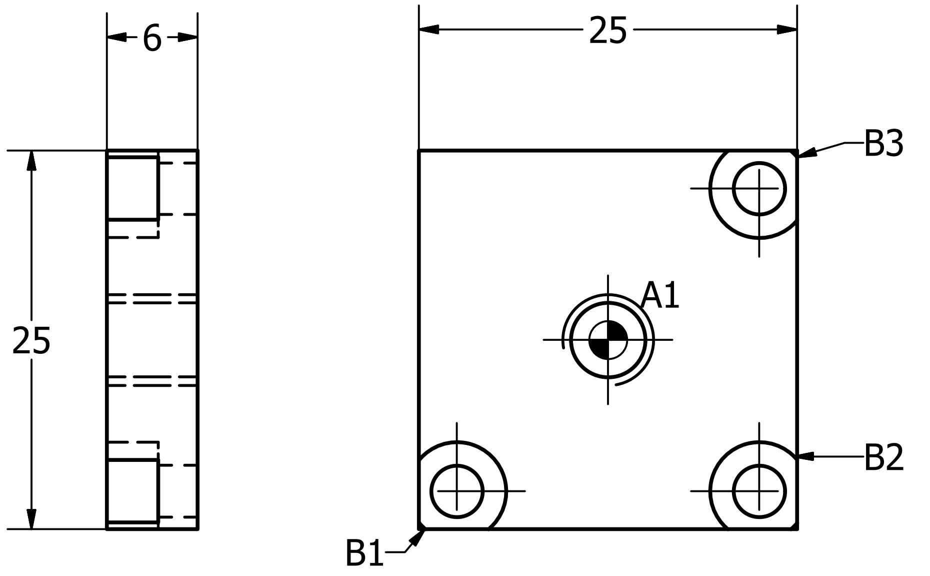

| HOLE TABLE | | | |
|---|---|---|---|
| HOLE | XDIM | YDIM | DESCRIPTION |
| A1 | 0 | 0 | M6x1 - 6H |
| B1 | -10 | -10 | Ø3,4 THRU<br>⌴ Ø6,5 ↧ 3,4 |
| B2 | 10 | -10 | |
| B3 | 10 | 10 | |

drive to dovetail adapter ( 2 : 1 )
1 off
Aluminium

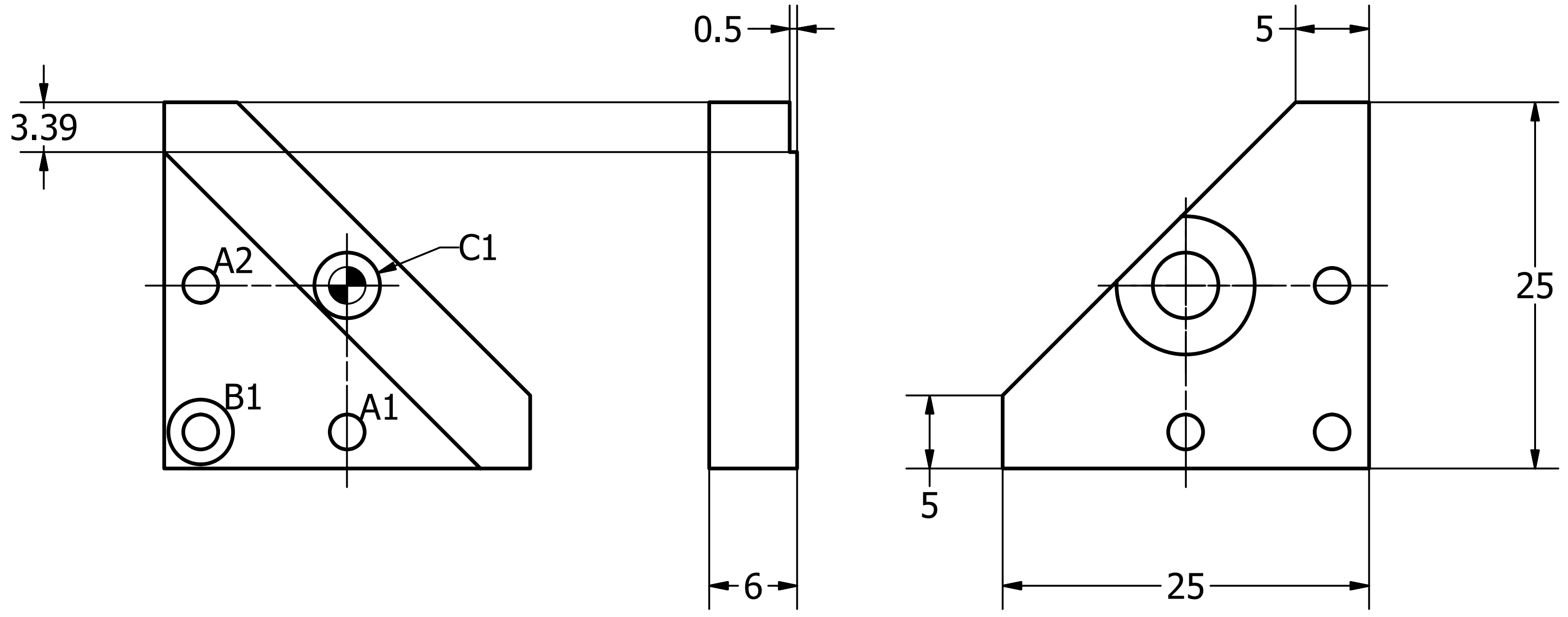

| HOLE TABLE | | | |
|---|---|---|---|
| HOLE | XDIM | YDIM | DESCRIPTION |
| A1 | 0 | -10 | Ø2,4 THRU |
| A2 | -10 | 0 | |
| B1 | -10 | -10 | Ø2,4 THRU<br>⌴ Ø4,4 ↧ 2,3 |
| C1 | 0 | 0 | Ø4,5 -8 DEEP<br>⌵ Ø9,4 X 90° |

mirror to drive adapter ( 2 : 1 )
1 off
Aluminium

ALUMINIUM
linear dimensions in mm

| DESIGNED BY | CHECKED BY | APPROVED BY | DATE | | DATE |
|---|---|---|---|---|---|
| map | | | | | 13/05/2019 |

| | ISSUE | SHEET |
|---|---|---|
| B24 Objective mount | 2 | 10 / 12 |

lifter plate ( 2 : 1 )
1 off
Aluminium

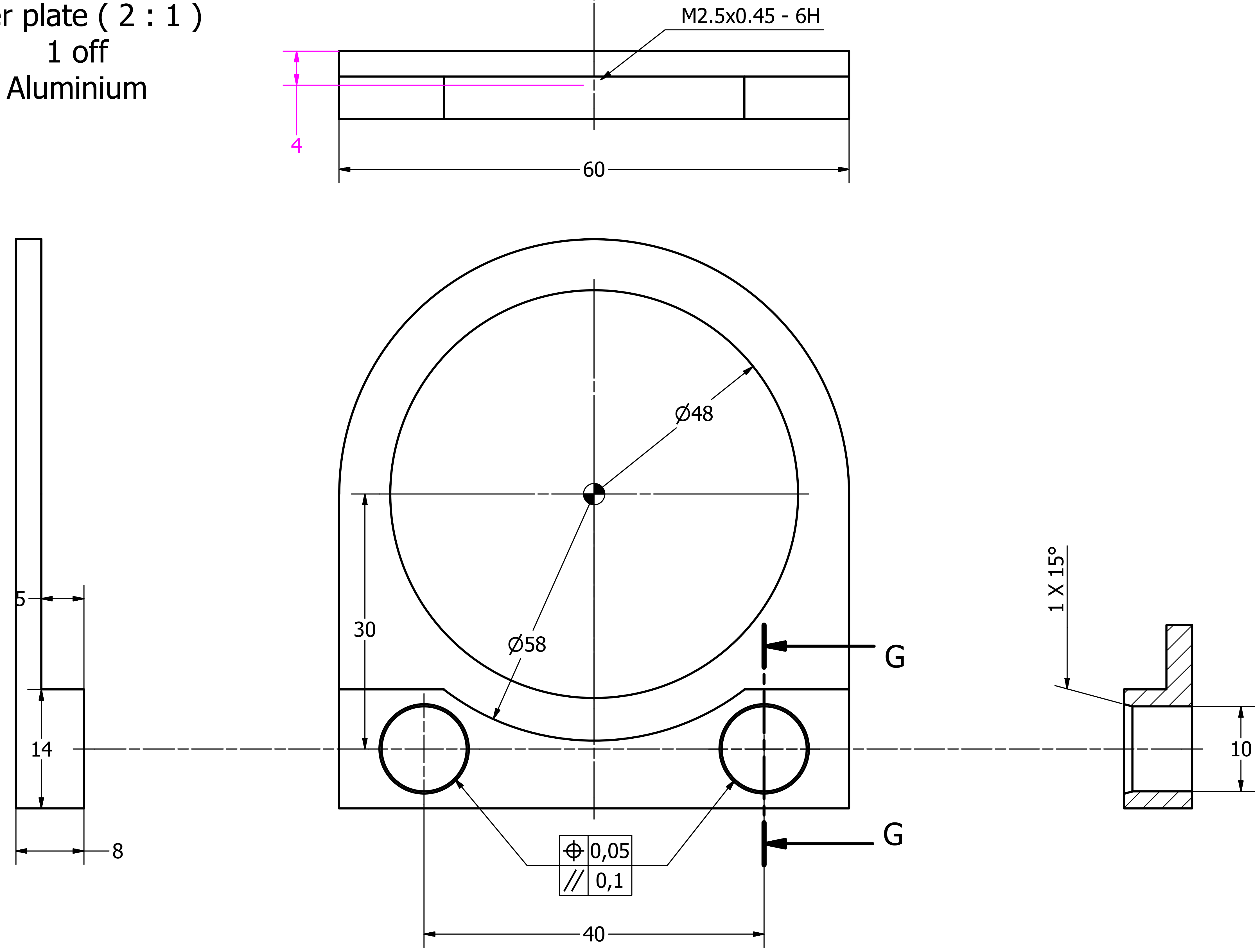

ALUMINIUM

linear dimensions in mm

| DESIGNED BY | CHECKED BY | APPROVED BY | DATE | | DATE |
|---|---|---|---|---|---|
| map | | | | | 13/05/2019 |

| | ISSUE | SHEET |
|---|---|---|
| B24 Objective mount | 2 | 11 / 12 |

#### objective plate ( 2 : 1 )
1 off
Aluminium

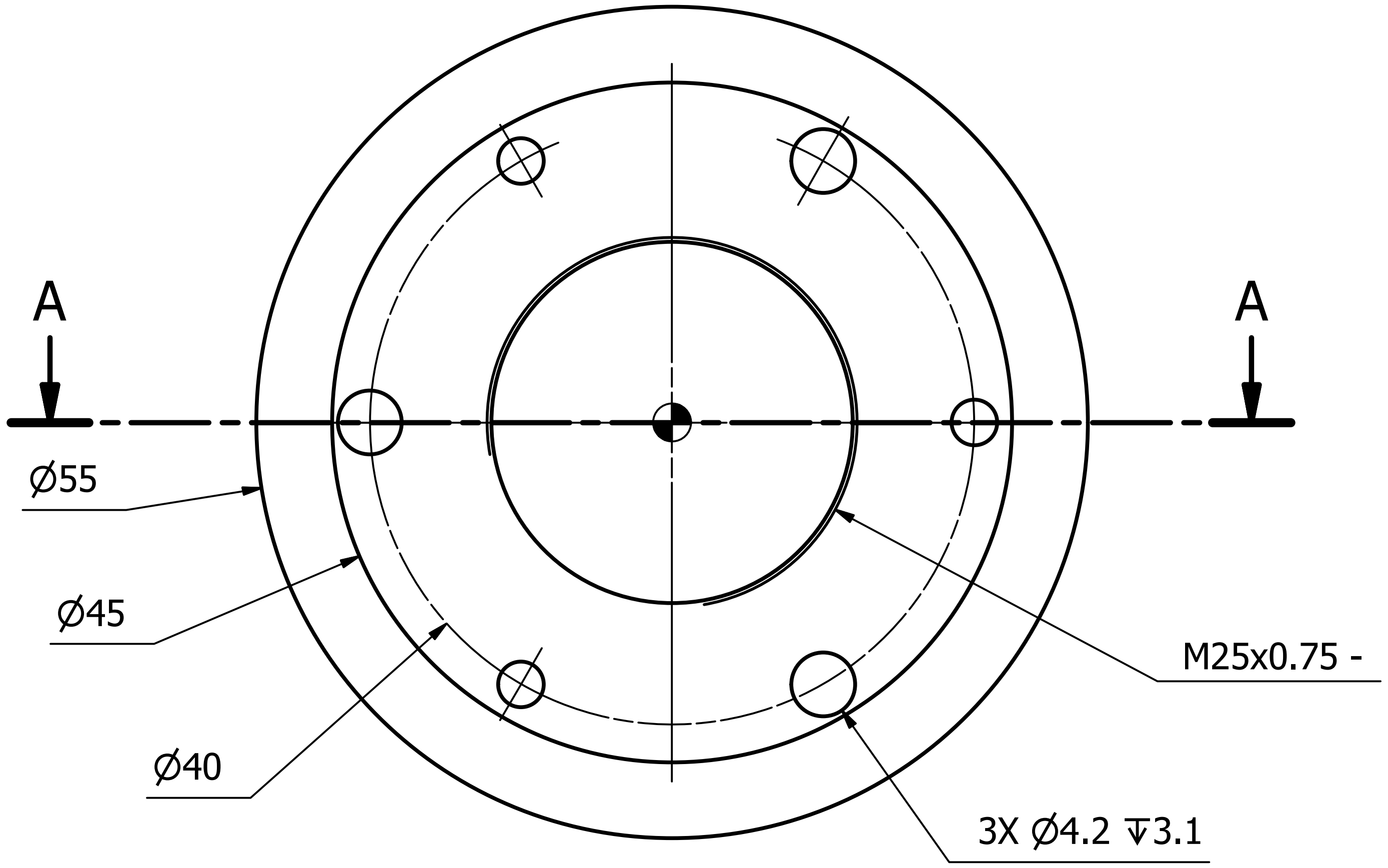

A-A ( 2 : 1 )

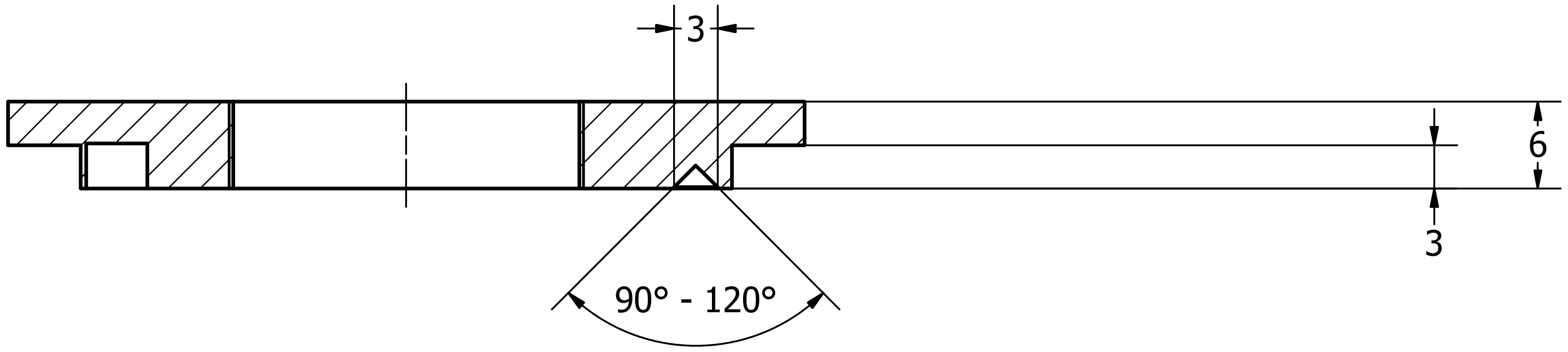

#### modified RC1 ( 2 : 1 )
1 part modification
Part supplied in aluminium

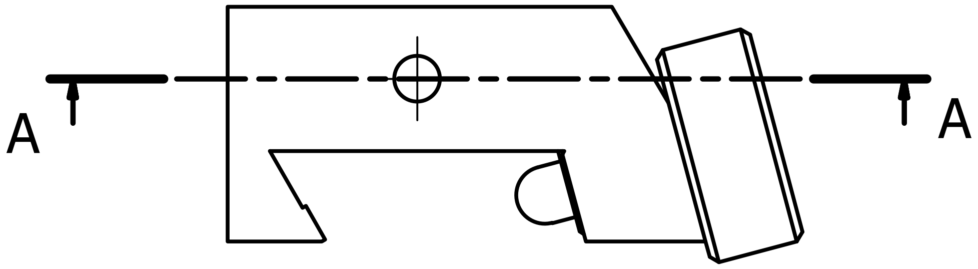

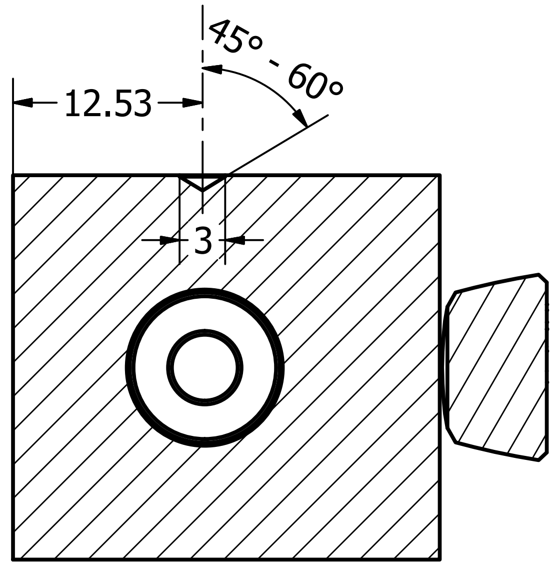

A-A ( 2 : 1 )

ALUMINIUM
linear dimensions in mm

| DESIGNED BY | CHECKED BY | APPROVED BY | DATE | | DATE |
|---|---|---|---|---|---|
| map | | | | | 13/05/2019 |

| | ISSUE | SHEET |
|---|---|---|
| B24 Objective mount | 2 | 12 / 12 |
