## Supplementary material for "CryoSIM: super resolution 3D structured illumination cryogenic fluorescence microscopy for correlated ultra-structural imaging": Custom parts diagrams: Plate.PiezoToLeadscrew.pdf

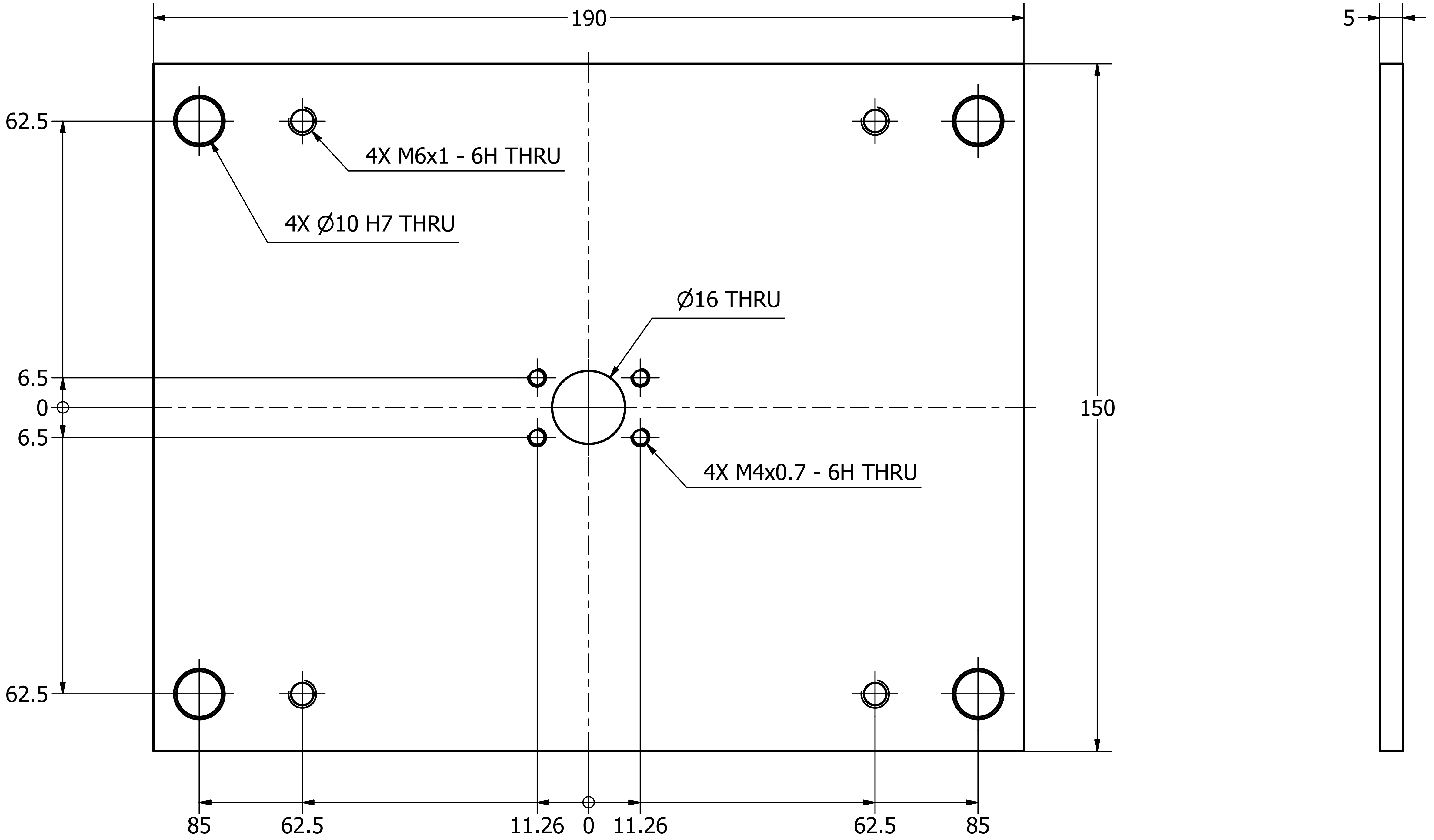


1 OFF, ALUMINIUM

| Designed by | Checked by | Approved by | Date | Date |
| --- | --- | --- | --- | --- |
| map | | | | 14/08/2018 |

3rd ANGLE PROJECTION

Micron Oxford

| Title | Edition | Sheet |
| --- | --- | --- |
| Plate.PiezoToLeadscrew | | 1 / 1 |
