## Supplementary material for "CryoSIM: super resolution 3D structured illumination cryogenic fluorescence microscopy for correlated ultra-structural imaging": Custom parts diagrams: RollerPlungerMount.pdf

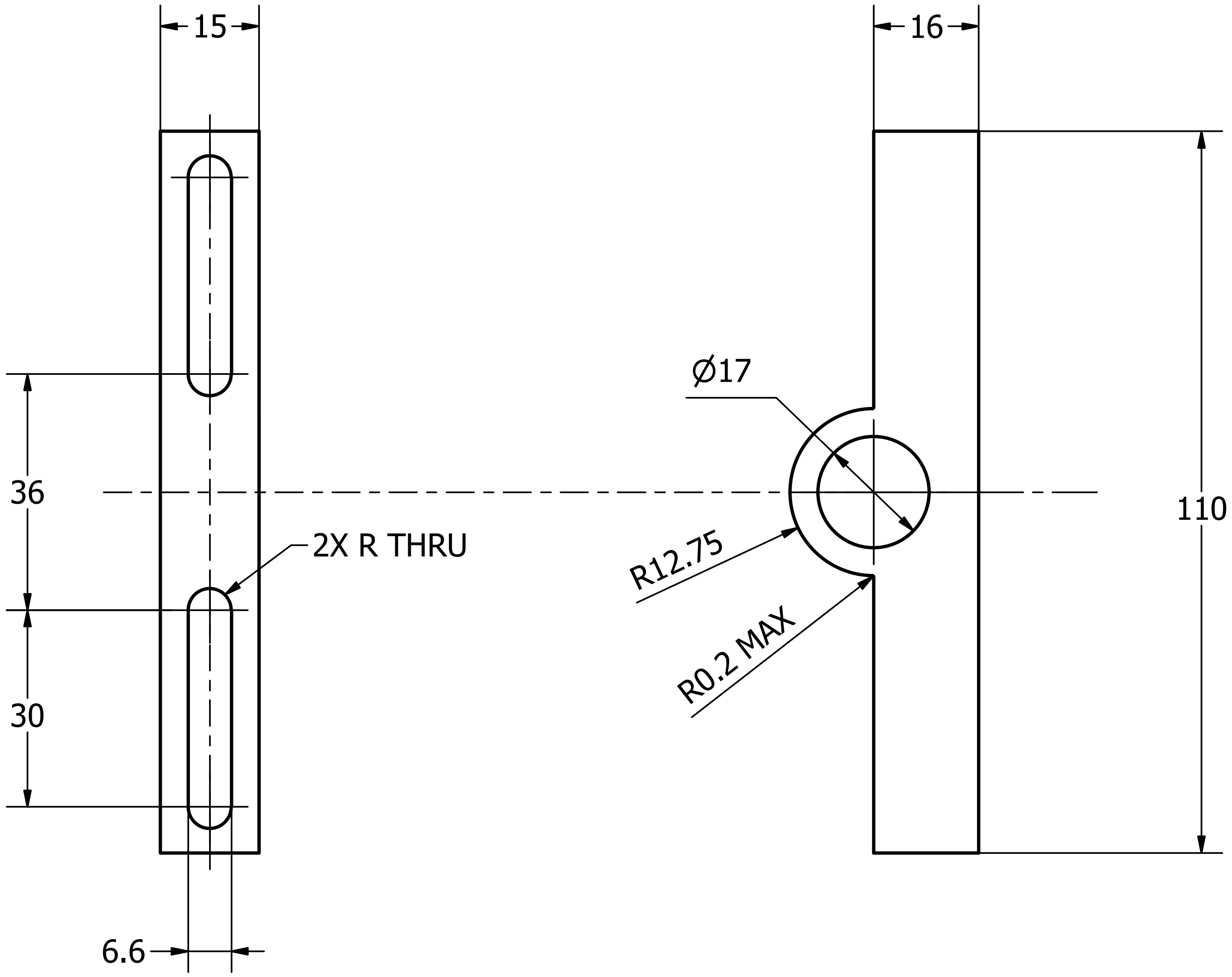


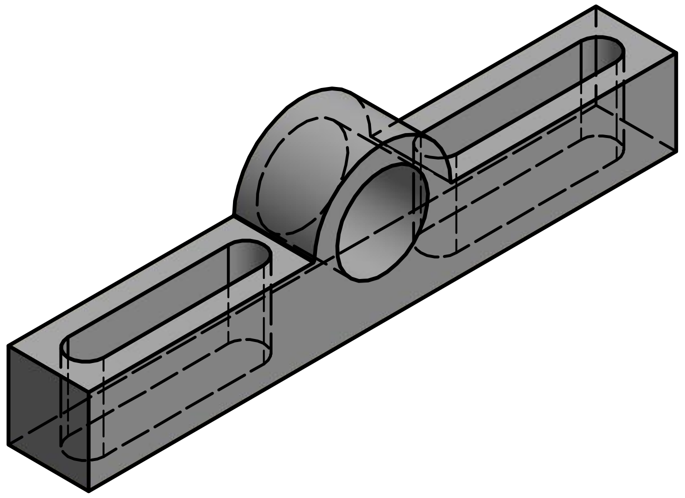

2 OFF, ALUMINIUM

| Designed by | Checked by | Approved by | Date | Date |
|---|---|---|---|---|
| map | | | | 06/06/2019 |

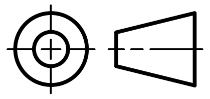


Micron Oxford

| | Edition | Sheet |
|---|---|---|
| RollerPlungerMount | | 1 / 1 |
